# VCBench: A Multi-Dimensional Benchmark for Single-Cell Foundation Models

**DOI:** 10.64898/2026.06.18.733146

**Authors:** L. Weidener, M. Brkić, M. Jovanović, E. Ulgac, A. Meduri

## Abstract

Single-cell foundation models are increasingly positioned as virtual cells, yet their capabilities are assessed by fragmented, largely single-task benchmarks that obscure where these models improve on simple baselines. VCBench addresses this by synthesizing four independent virtual-cell frameworks into seven capability dimensions: perturbation response prediction, cross-species universality, gene regulatory network (GRN) inference, modality integration, temporal dynamics, multi-scale integration, and in silico experimentation. Each dimension is assessed for operational testability under current architectures and datasets: five admit direct or proxy evaluation, while multi-scale integration and in silico experimentation are structurally untestable as end-to-end tasks. We evaluate five foundation models (Geneformer, scGPT, UCE, TranscriptFormer, Arc State) against pre-registered linear and nearest-neighbor baselines across the five testable dimensions, and report three findings. First, the baselines match or exceed every foundation model on four of the five scored dimensions, replicating the reported competitiveness of linear baselines on perturbation prediction and extending it to cross-species transfer, GRN inference, and temporal ordering. Second, TranscriptFormer alone exceeds the strongest baseline on cross-modal RNA-to-protein prediction (53% Pearson improvement, with a documented contamination caveat) and is the only model to reach Level 2 in the pre-registered Virtual Cell (VC) Level rubric; the architectural choice behind this advantage simultaneously causes a spectral collapse that destroys its temporal-ordering performance, a tradeoff invisible to single-task benchmarks. Third, no foundation model publishes a complete cell-level training manifest, leaving data contamination undetectable to users. Alongside the benchmark, VCBench releases a Contamination Reporting Schema and contributes two further methodological tools: a common-label-set protocol that controls for class-count confounds in cross-species transfer, and a spread-error correlation probe for epistemic calibration.

## 1 Background

A computational model that faithfully simulates cellular behavior would transform biology: predicting cellular responses to genetic perturbation, forecasting differentiation trajectories, identifying drug targets, and reducing animal experimentation in preclinical safety assessment [2]. Several single-cell foundation models are positioned as steps toward this vision, sharing the transformer backbone but differing in tokenization, pretraining objective, and training corpus. Geneformer [3] uses rank-value gene tokenization with masked learning on 30 million cells from Genecorpus-30M. scGPT [4] extends this with generative pretraining on 33 million CELLxGENE cells and a uni-fied token scheme for multi-omic data. Universal Cell Embeddings (UCE) [5] tokenizes genes through ESM-2 protein-language-model features, producing species-agnostic embeddings. TranscriptFormer [6] scales the autoregressive approach to 112 million cells across 12 species. Arc State [7] departs from single-cell tokenization, operating on sets of cells through a state-embedding module trained on ~167 million observational cells and a state-transition module trained on 100 million perturbed observations.

Four independent frameworks articulated what a virtual cell should achieve. Bunne et al. [2] described a multi-scale, multi-modal AI Virtual Cell. Roohani et al. [8] operationalized perturbation response into the Virtual Cell Challenge, framed as “a Turing test for the virtual cell.” Noutahi et al. [9] proposed a Predict-Explain-Discover framework structured around three capability tiers. Dibaeinia et al. [10] formalized perturbation prediction as a causal transport problem and argued current architectures are fundamentally misaligned with the task.

Recent empirical benchmarks increase the urgency. Ahlmann-Eltze et al. [1] showed that simple linear baselines match or exceed five foundation models on perturbation prediction; Kedzierska et al. [11] demonstrated comparable failures on zeroshot cell-type embedding; Kendiukhov [12] found trivial gene-level baselines outperform attention- and correlation-derived regulatory edges in Geneformer and scGPT, with attention contributing no incremental signal to perturbation prediction. It remains unclear how current foundation models perform across the capability dimensions envisioned for virtual cells. Existing benchmarks evaluate perturbation prediction extensively but do not test whether models generalize across species, infer regulatory networks, simulate temporal dynamics, or integrate across scales.

This research aims to consolidate the capabilities that these frameworks envision for a virtual cell into a single benchmark and to measure where current foundation models meet them. A structured synthesis of the four frameworks identifies seven recurring capability dimensions, and each is assessed for operational testability under current models and datasets: three admit direct evaluation (perturbation prediction, cross-species universality, gene regulatory network (GRN) inference), two admit proxy tasks (modality integration via cellular indexing of transcriptomes and epitopes by sequencing (CITE-seq), and temporal dynamics via time-course ordering), and two (multi-scale integration, in silico experimentation) are structurally untestable as end-to-end tasks with current models and datasets. The five testable dimensions are scored for each foundation model in a capability matrix that distinguishes the regimes where simple baselines remain competitive from those where foundation models add architectural value. Alongside the empirical evaluation, VCBench contributes three reusable methodological tools introduced in the course of construction: a common-label-set evaluation protocol that controls for class-count confounds in cross-species transfer, a spread-error correlation probe for epistemic calibration, and a contamination-reporting schema that addresses the transparency deficit surfaced by the audit.

## 2 Methods

Pre-submission methodological audits, protocol amendments, and operational caveats are documented chronologically in Supplementary Note 2; key items affecting interpretation of Table 2 are also flagged inline in the relevant Methods subsection below. Code paths for each scored dimension were independently verified against their Methods descriptions; verification logs are released alongside the benchmark code.

### 2.1 Framework synthesis and operational testability

A structured review of four virtual-cell frameworks (Bunne et al. [2], Roohani et al. [8], Noutahi et al. [9], Dibaeinia et al. [10]) identifies seven recurring capability dimensions: (A) perturbation response prediction; (B) cross-species universality; (C) gene regulatory network inference; (D) modality integration; (E) temporal dynamics; (F) multi-scale integration; and (G) in silico experimentation (Table 1). Each dimension is then assessed for operational testability under current model architectures and available datasets, yielding three categories: directly testable (A, B, C); testable via proxy tasks (D as cross-modal CITE-seq prediction; E as time-course ordering); structurally untestable as end-to-end tasks (F, G). Figure 1 visualizes the framework synthesis, the evaluation architecture (5 foundation models x5 scored dimensions with regime per cell), the dataset-dimension map, and the pre-registered Virtual Cell Level thresholds adopted from Noutahi et al. [9].

**Table 1.**
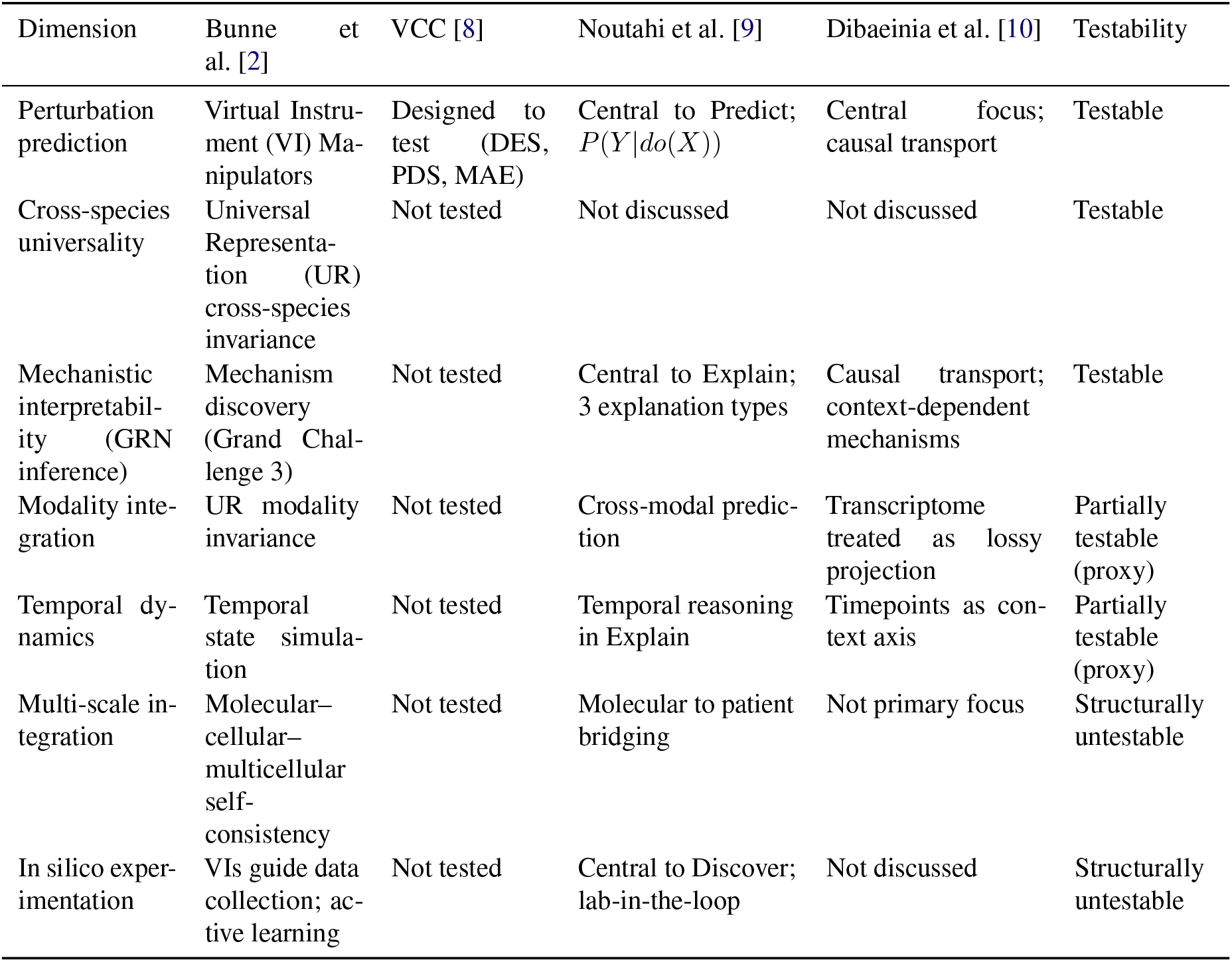
Virtual cell capability dimensions across four independent frameworks.

**Figure 1.**
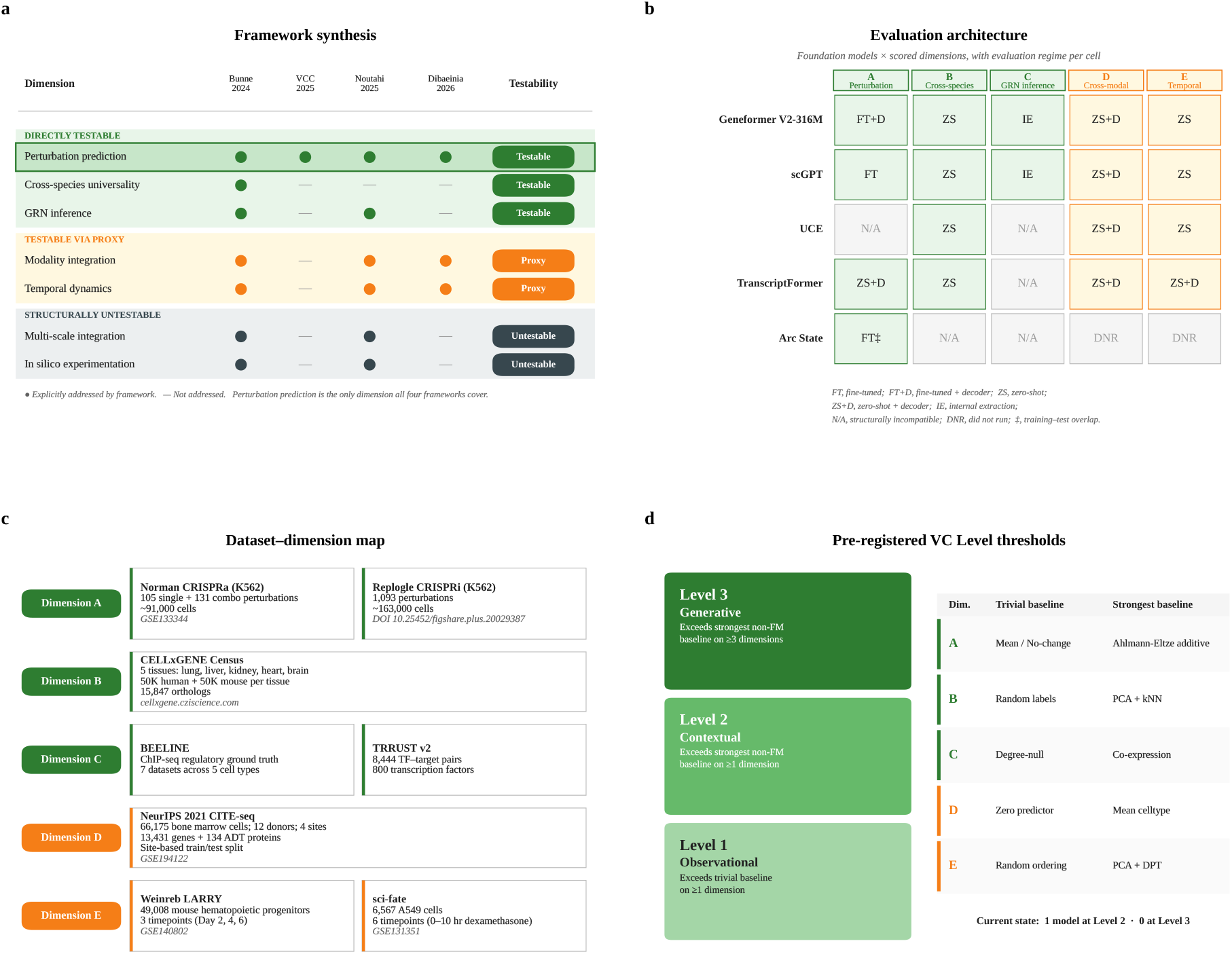
Framework synthesis and evaluation architecture. (a) Four independent virtual-cell frameworks synthesized into seven capability dimensions. (b) Evaluation architecture: 5 foundation models ×5 scored dimensions with regime per cell (FT, FT+D, ZS+D, ZS, IE). (c) Dataset-dimension map. (d) Pre-registered Virtual Cell Level thresholds adopted from Noutahi et al. [9].

### 2.2 Datasets and preprocessing

Norman CRISPRa [13]: 105 single-gene and 131 combination perturbations in K562 cells, 91,000 cells total (GSE133344). GEARS test split [14] (107 perturbations) is used; the 71-perturbation additive-evaluable subset and 36-perturbation novel subset are evaluated separately. CELLxGENE Census [15]: cross-species cell-type transfer corpus drawn from 5 tissues (lung, liver, kidney, heart, brain), 50,000 human and 50,000 mouse cells per tissue, restricted to 15,847 protein-coding orthologs from the Ensembl one-to-one mapping (cellxgene.cziscience.com); see Breschi et al. [16] for human–mouse comparative-transcriptomics context.

BEELINE [17]: 7 ChIP-seq-grounded GRN datasets across 5 cell types (hESC, hHEP, mDC, mESC, mHSC; human/mouse embryonic stem, hepatocyte, dendritic, and hematopoietic stem cells). TRRUST v2 [18]: 8,444 curated transcription factor (TF)–target pairs covering 800 transcription factors.

NeurIPS 2021 CITE-seq [19]: 66,175 bone marrow cells; 12 donors; 4 sites; 13,431 genes and 134 surface proteins (GSE194122). Site-based train/test split.

Weinreb LARRY [20]: 49,008 mouse hematopoietic progenitors with 3 timepoints (days 2, 4, and 6) (GSE140802). sci-fate [21]: 6,567 A549 cells with 6 timepoints across 0–10 h dexamethasone treatment (GSE131351).

### 2.3 Model selection and configurations

Five foundation models are evaluated: Geneformer V2-316M [3]; scGPT [4]; UCE [5]; Transcript-Former [6]; Arc State [7] (state-embedding + state-transition module composition). Each model is evaluated using the strongest protocol available to it (fine-tuning where supported on Dimension A; zero-shot embedding probes on Dimensions B, D, E; internal-state extraction on Dimension C). The capability matrix (Table 2) annotates each entry with its evaluation regime: FT (fine-tuned), FT+D (fine-tuned with decoder), ZS (zero-shot), ZS+D (zero-shot with decoder), IE (internal extraction). Cells marked N/A indicate the model architecturally cannot produce the required output type for that dimension; cells marked DNR (did not run) indicate the evaluation was attempted but did not produce valid results due to runtime errors, incompatible data formats, or absence of a public evaluation interface.

**Table 2.** Capability matrix for five single-cell foundation models across five evaluation dimensions. Each cell reports the primary metric for the given model-dimension combination with evaluation regime in parentheses (FT, fine-tuned; FT+D, fine-tuned with decoder; ZS, zero-shot; ZS+D, zero-shot with decoder; IE, internal extraction). Cells exceeding the strongest non-foundation-model baseline are bolded. Dimension B values are aggregated under the native-label protocol; under the common-label-set protocol no foundation model exceeds the PCA+kNN baseline at the aggregate level, though TranscriptFormer marginally exceeds the baseline on liver per-tissue (Methods, Supplementary Table 2). Secondary metrics (DES, weighted F1, RMSE, balanced accuracy) are reported in Supplementary Tables 2, 3 and Methods. ^*†*^ TranscriptFormer Weinreb reported as bootstrap mean over 10 random 5,000-cell subsamples (std 0.078). ^*‡*^ Arc State Norman result reported under the corrected configuration.

| Model | A: PRR | B: MacroF1 | C: AUROC | C: AUPRC | C: EPR | D: Pearson | E: τ-b |
| --- | --- | --- | --- | --- | --- | --- | --- |
| Geneformer V2-316M | 0.627 (FT+D) | 0.181 (ZS) | 0.626 (IE) | 0.001 (IE) | 0.000 (IE) | 0.001 (ZS+D) | -0.017 (ZS) |
| scGPT (fine-tuned) | 0.503 (FT) | 0.012 (ZS) | 0.519 (IE) | 0.003 (IE) | **20.05** (IE) | 0.064 (ZS+D) | -0.057 (ZS) |
| UCE 33-layer | N/A | 0.027 (ZS) | N/A | N/A | N/A | 0.132 (ZS+D) | 0.136 (ZS) |
| TranscriptFormer | -0.174 (ZS+D) | 0.156 (ZS) | N/A | N/A | N/A | **0.232** (ZS+D) | 0.046<sup>†</sup> |
| Arc State | 0.402<sup>‡</sup> (FT) | N/A | N/A | N/A | N/A | DNR | DNR |
| Additive baseline | 0.890 | — | — | — | — | — | — |
| Mean baseline | 0.579 | — | — | — | — | 0.152 | — |
| No-change baseline | 0.000 | — | — | — | — | — | — |
| PCA+kNN | — | 0.166 | — | — | — | — | — |
| Co-expression | — | — | 0.558 | 0.004 | 15.50 | — | — |
| Degree-null | — | — | 0.500 | 0.0003 | 1.13 | — | — |
| pySCENIC | — | — | 0.501 | 0.0011 | 3.50 | — | — |
| scLinear | — | — | — | — | — | 0.129 | — |
| Mean celltype | — | — | — | — | — | 0.152 | — |
| PCA+ridge | — | — | — | — | — | 0.110 | — |
| PCA+DPT | — | — | — | — | — | — | 0.190 |

#### Geneformer V2-316M

316M-parameter masked-language-model variant, fine-tuned on Norman with token-deletion perturbation and a learned decoder head; zero-shot embeddings extracted from the pretrained checkpoint for Dimensions B, D, E; gene-attention extraction for Dimension C.

#### scGPT

pretrained on 33M cells; fine-tuned on Norman; zero-shot embeddings for Dimensions B, D, E; gene-embedding cosine similarity for Dimension C.

#### UCE

pretrained ESM-2-tokenized representations; zero-shot evaluation across all evaluable dimensions; N/A on Dimensions A and C (no native fine-tuning interface; no gene-level edge prediction).

#### TranscriptFormer

autoregressive multi-species pretraining; ZS+D protocol on all evaluable dimensions. N/A on Dimension C (the public release lacks a gene-attention extraction interface compatible with the BEELINE/TRRUST evaluation pipeline; see Supplementary Note 2 §S2.1).

#### Arc State

state-embedding module for cross-modal probes; state-transition module for perturbation prediction. Fine-tuned on Norman under a corrected GEARS split (an earlier configuration we wrote contained a train-test leak; see Methodological note below). N/A or DNR on Dimensions B, C, D, E reflecting architectural incompatibility (set-level rather than cell-level outputs) or absence of a public evaluation interface.

Stack [22] (in-context learning of single-cell biology) and scPRINT [23] (denoising-based GRN inference) were considered but excluded as architecturally incompatible with VCBench’s held-out evaluation protocol or narrower in scope than the multi-dimensional capability profile evaluated here.

### 2.4 Evaluation protocols

#### Dimension A (perturbation response prediction)

Norman CRISPRa [13] serves as the primary benchmark; the 107-perturbation GEARS test split [14] is decomposed into a 71-perturbation additive-evaluable subset (both single-gene components observed in training) and a 36-perturbation novel subset (at least one component held out). The Perturbation Response Recovery (PRR) score is the mean per-perturbation Pearson correlation between predicted and observed log-fold-change (LFC) vectors relative to control:

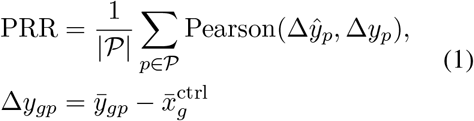

where *P* is the set of test perturbations, *g* indexes genes, 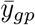 is the mean expression of gene *g* across cells receiving perturbation *p*, and 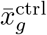 is the mean control expression; predicted quantities are defined analogously. PRR is bounded in [ −1, 1]. We adopt PRR in lieu of Cell-Eval’s canonical Perturbation Discrimination Score (which requires a closed candidate-perturbation pool not compatible with the held-out single-perturbation evaluation setting used here); PRR uses the same per-perturbation Δ-expression construction but summarizes via correlation rather than ranking. Note that Figure 3 carries “PDS” axis and annotation labels from an earlier figure-generation pass; the values shown are identical to the renamed PRR. The Direction Score (DES; sign-agreement on the top-20 most-perturbed genes per perturbation, formal definition in Eq. 3) and mean absolute error (MAE) are reported as secondary metrics. The Ahlmann-Eltze additive baseline (Eq. 2) is the strongest non-foundation-model baseline; the no-change baseline (zero predicted change) is the trivial baseline. Mathematical definitions for all remaining metrics, baselines, and statistical procedures are consolidated in §Mathematical definitions.

#### Dimension B (cross-species universality)

Each model’s cross-species transfer score is computed under two protocols. The native-label protocol scores each model on the cell-type vocabulary that survives its own tokenizer and filter, which differs across models and produces non-comparable label sets. The common-label-set protocol restricts every model and the strongest baseline to the same cell-type vocabulary per tissue (Eq. 4), removing the class-count confound that previously inflated foundation-model performance under the native protocol. PCA+kNN (k=5, cosine metric) is the strongest baseline; random label assignment is the trivial baseline. Macro F1 (Eq. 5) is the primary metric; weighted F1 is reported as secondary. The common-label-set protocol is the binding protocol for VC Level assignments and bolding decisions; native-protocol values are reported in Table 2 for direct cross-paper comparability with the original Census-derived numbers [15], with full per-tissue breakdowns under both protocols in Supplementary Table 2.

#### Dimension C (gene regulatory network inference)

Area under the precision-recall curve (AUPRC) and Early Precision Ratio (EPR; Eq. 6) are jointly required to exceed all three baselines (co-expression, degree-null, pySCENIC [38]) for a passing claim. Area Under the Receiver Operating Characteristic curve (AUROC) is additionally reported as a ranking diagnostic but is not part of the passing rule. Bootstrap-resampled 95% confidence intervals (CIs) on AUPRC are computed via 1,000 paired iterations (model vs. baseline on the same edge set; Eq. 7). Co-expression is the strongest baseline; degree-null is the trivial baseline.

#### Methodological note: GRN passing rule amendment

The pre-registered passing rule originally required AUPRC alone to exceed the strongest baseline; after initial results showed scGPT EPR substantially exceeds all baselines while AUPRC remains at the noise floor, the rule was amended to the conjunctive AUPRC + EPR criterion. This amendment is disclosed before any model is reported as passing or failing. The amended rule is more conservative than the original (no model passes under either rule, but the conjunctive rule prevents future models from passing on EPR alone with poor AUPRC).

#### Dimension D (modality integration; cross-modal CITE-seq prediction)

RNA embeddings are extracted from each foundation model on the NeurIPS 2021 CITE-seq dataset [19]; a ridge regression decoder is fit to predict surface-protein abundance (134 antibody-derived tags, ADTs) from the RNA embedding, using a site-based train/test split. Per-protein Pearson correlation is the primary metric; root-mean-square error (RMSE) is the secondary metric. Mean-celltype prediction (Eq. 8) is the strongest baseline; scLinear [39] and PCA+ridge are reported as additional reference baselines (Table 2); the zero predictor is the trivial baseline. The contamination caveat (GSE194122 confirmed in CELLxGENE Census) is documented in the contamination Results sub-section and Supplementary Note 2.

#### Dimension E (temporal dynamics; time-course ordering). Diffusion pseudotime (DPT)

[40] is computed on each foundation model’s embedding of the relevant time-course dataset (Weinreb LARRY [20], sci-fate [21]). Kendall’s τ-b (Eq. 9) between predicted pseudotime and ground-truth time annotation is the primary metric; balanced accuracy across timepoints is the secondary metric. PCA+DPT (50 components, default DPT parameters) is the strongest baseline; random ordering is the trivial baseline. For TranscriptFormer on Weinreb, full-dataset DPT eigensolution failed due to near-duplicate embeddings producing a degenerate graph Laplacian (ARPACK eigensolver non-convergence; Figure 7 documents the spectral structure). The TranscriptFormer Weinreb result is computed via 10-fold bootstrap on 5,000-cell subsamples (mean *±* standard deviation).

### 2.5 Mathematical definitions

This subsection consolidates formal definitions for every metric, baseline, and statistical procedure used in VCBench. Notation: *i* indexes cells, *g* indexes genes, *p* indexes perturbations, *c* indexes cell types, *t* indexes tissues, and *m* indexes evaluated methods (foundation models and baselines combined). *x*_*ig*_ denotes observed expression, 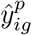 predicted expression under perturbation *p*, and 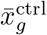 the mean control expression of gene *g*. The PRR equation is given in §Methods (Eq. 1).

#### Additive baseline (Dimension A)

For a combinatorial perturbation *A* + *B* in which both single-gene components *A* and *B* are observed as singletons in training, the Ahlmann-Eltze additive prediction is

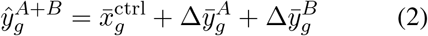

with 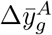 and 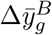 the observed mean LFCs for the singleton perturbations. The baseline is structurally inapplicable when either component is held out, defining the 71-perturbation additive-evaluable subset and 36-perturbation novel subset of the 107-perturbation Norman GEARS test split.

#### Direction Score (Dimension A, secondary)

For each test perturbation *p*, let 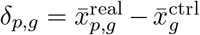 and 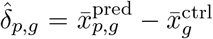 denote the observed and predicted pseudo-bulk Δ-expression vectors. With *T*_*K*_(*p*) the indices of the top *K* = min(20, *G*) genes ranked by |*δ*_*p,g*_|, the per-perturbation direction score is the fraction of those genes on which predicted and observed Δ agree in sign:

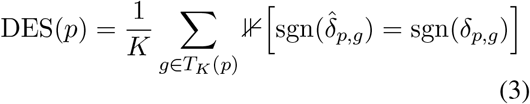

and the dataset-level score is 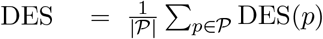. The *K* = 20 cap and pseudo-bulk averaging are pipeline-level choices fixed at evaluation time; the additive-baseline DES values reported in Cell-Eval canonical mode [7] use a different aggregation and are not directly comparable to the FM DES values reported here, paralleling the PRR/PDS distinction discussed above.

#### Common-label-set protocol (Dimension B)

For each tissue *t*, let 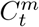 denote the cell-type vocabulary admitted by method *m*. The commonlabel-set protocol restricts evaluation per tissue to

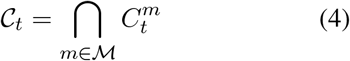

the intersection over all evaluated methods *ℳ* in tissue *t*. Cells whose annotation lies outside *C*_*t*_ are excluded from scoring under this protocol. The common-label-set protocol is the binding protocol for VC Level assignments and bolding decisions.

#### Macro and weighted F1 (Dimension B)

For a label set *C* and class-conditional F1 scores *F* 1_*c*_,

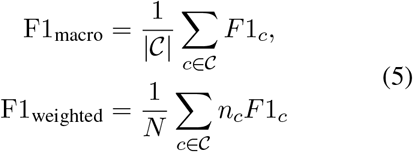

where *n*_*c*_ is the support of class *c* and *N* = Σ_*c*_ *n*_*c*_. Macro F1 is the primary metric. Under the native protocol the macro F1 of method *m* depends mechanically on 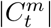, producing the class-count confound that the common-label-set protocol corrects.

#### Early Precision Ratio (Dimension C)

Let *K* be the number of true positive edges in the ground truth (so that *K/N* is the random-baseline expected precision over *N* candidate edges) and let Precision@K denote the precision of the model’s top-*K* predicted edges, evaluated at the same *K*. The early precision ratio normalizes top-*K* precision against the random-baseline expectation:

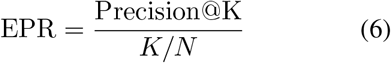

EPR = 1 corresponds to random ranking; EPR *>* 1 indicates true edges are concentrated above chance among the top-*K* predictions. The conjunctive passing rule for Dimension C requires both AUPRC and EPR to exceed all three baselines.

#### Bootstrap confidence intervals and paired tests (Dimension C)

For each foundation model *m* and baseline *b, B* = 1,000 paired bootstrap iterations indexed *t* = 1, …, *B* resample the evaluation edge set *E* with replacement, computing 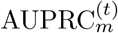 and 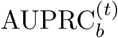 on the same resam-pled edges per iteration. The 95% CI is the [2.5th, 97.5th] percentile of the bootstrap distribution. The paired one-sided p-value is

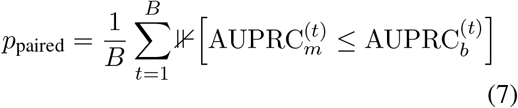

with Benjamini-Hochberg FDR correction across the model ×baseline pairwise tests.

#### Mean-celltype baseline (Dimension D)

For each test cell *i*, let *ĉ*(*i*) denote the cell type predicted by a cosine-distance kNN classifier (*k* = 5) on the training-site RNA embeddings. For each surface protein π ∈ {1, …, 134}, the mean-celltype prediction is

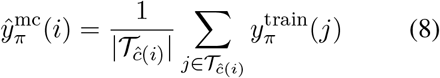

with *T*_*c*_ the set of training cells annotated to cell type *c*. The baseline thus carries no information beyond the predicted cell type identity.

#### Kendall’s *τ* -b (Dimension E)

For predicted and ground-truth orderings on *n* cells with *n*_*c*_ concordant and *n*_*d*_ discordant pairs,

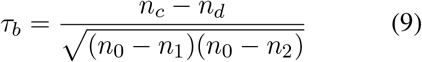

with 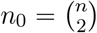 and *n*_1_, *n*_2_ the tie corrections on the predicted and ground-truth orderings respectively. The *τ*-b form is required because timecourse ground-truth timepoints are discrete and produce many ties. Per-dataset *τ*_*b*_ values are aggregated across sci-fate and Weinreb LARRY by unweighted arithmetic mean.

#### Spread-error correlation probe (epistemic calibration)

For each test perturbation *p*, the spread is the variance of the predicted LFC vector across genes, and the error is the per-perturbation MAE against ground truth:

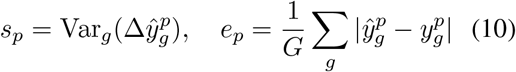

The spread-error correlation is 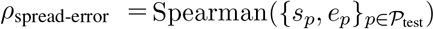. Positive ρ indicates the model assigns higher spread to harder predictions; ρ ≈ 0 indicates spread carries no information about prediction quality. The probe is necessary but not sufficient for epistemic calibration: passing requires distributional output, ensembles, MC dropout, or conformal wrappers, none of which the tested models provide.

### 2.6 Methodological notes

#### Methodological note: train-test leak in our Arc State evaluation

During VCBench development, an evaluation of Arc State on the Norman CRISPRa dataset that we performed using a Norman-specific TOML configuration file `(norman_fewshot.toml)`, which we constructed from Arc Institute’s published `examples/fewshot.toml` template, produced an inflated PRR of 0.949 via a train-test leak: the cell-type filter we wrote `([zeroshot] “norman.double_perts” = “test”)` silently matched zero rows in the K562-only Norman dataset, because the State framework does not validate that cell-type filter values match any cells in the loaded data. All 107 nominally held-out test perturbations therefore remained in the training pool. The leak was identified during pre-submission audit; we retrained Arc State under the correct GEARS split `(norman_gears_ split.toml,` with explicit train/test perturbation lists matching the canonical GEARS simulation split (seed 1): 138 training perturbations plus the non-targeting control and 107 held-out test perturbations, with zero overlap) for the reported PRR 0.402 result. Training loss converged to 0.027 while validation loss oscillated and diverged to approximately 0.40, indicating overfitting to the 138-perturbation training set. A one-line assertion at config-load time (raise if a cell-type filter matches zero rows in the loaded dataset) would have caught this configuration error at training-launch and is recommended for future versions of the State framework. Full audit narrative including the three independent diagnostic signatures and the evaluator anchor convention is in Supplementary Note 3 (Dimension A protocol, Arc State paragraph).

#### Methodological note: TranscriptFormer Dimension C N/A

TranscriptFormer is reported N/A on Dimension C rather than evaluated and scored. The model’s public release lacks a gene-attention extraction interface compatible with the BEELINE/TRRUST evaluation pipeline used for Geneformer and scGPT. Implementing such an interface would require either monkey-patching the released code or training a separate gene-relationship probe, neither of which preserves an apples-to-apples comparison. Full discussion in Supplementary Note 2 §S2.1.

#### Methodological note: Dimension B common-label-set evaluation

Dimension B is reported under the common-label-set protocol in which every model and the PCA+kNN baseline are restricted per tissue to the intersection of cell-type vocabularies admitted by all evaluated methods. This removes a class-count confound that arises when foundation models and baselines are scored against different cell-type vocabulary sizes. Under the common-label-set protocol, PCA+kNN aggregate macro F1 is 0.497 versus Geneformer 0.171, scGPT 0.123, and UCE 0.379 (all five tissues); TranscriptFormer’s lung+liver F1 is 0.351 versus PCA+kNN’s matched lung+liver value of 0.456. Per-tissue, PCA+kNN exceeds every foundation model on lung, heart, kidney, and brain; on liver, TranscriptFormer (macro F1 0.495) marginally exceeds PCA+kNN (0.446). Native-protocol numbers are reported alongside in Supplementary Table 2. TranscriptFormer’s common-set evaluation is restricted to lung and liver, the two tissues where TranscriptFormer holds species embeddings; full-tissue common-set evaluation is scoped to a future revision. Full audit narrative in Supplementary Note 2 §S2.3.

#### Methodological note: capability matrix numerical provenance

Numbers in the capability matrix (Table 2) are sourced from the canonical evaluation pipeline at the v1.0.0 release of the benchmark code (release tag and commit SHA in Data and code availability). Dimension A values use the GEARS partition definition matching `combo_seen0+1+2` versus `unseen_single` (71 shared / 36 novel out of 107 GEARS test perturbations). A pre-submission audit reconciled stale entries against the canonical pipeline; full reconcil-iation log is in Supplementary Note 2 §S2.2. Detailed per-dimension methodology including exact hyperparameters and library versions is in Supplementary Note 3.

### 2.7 VC Level mapping

Foundation models are assigned to one of four Virtual Cell (VC) Levels based on their per-dimension scores: Level 0 (no dimension exceeds trivial baseline); Level 1 (exceeds trivial baseline on at least one dimension); Level 2 (exceeds strongest non-foundation-model baseline on at least one dimension); Level 3 (exceeds strongest non-foundation-model baseline on three or more dimensions). The Level definitions and per-dimension thresholds are pre-registered before benchmark execution and adopted from Noutahi et al. [9] with minor adaptations for operational testability. Threshold values per dimension are listed in Figure 1d.

### 2.8 Use of large language models

Large language models (Anthropic Claude family) were used during manuscript preparation for editorial assistance: text revision, structural reorganization between drafts, and pre-submission consistency auditing (numerical cross-references, claimtable coherence, methodological-note ordering). All scientific content, results, code, and analyses were authored by the human authors. LLM outputs were independently reviewed and edited; no claims, numbers, or references in this manuscript originate solely from LLM generation without author verification.

## 3 Results

### 3.1 Pretraining data overlap is pervasive but structurally asymmetric

A systematic contamination audit (Supplementary Table 5) reveals structural asymmetry. No evaluated model publishes a complete cell-level training manifest, making exhaustive detection impossible. CELLxGENE Census (the primary source for scGPT, UCE, TranscriptFormer, and Arc State’s embedding model) explicitly excludes cell-culture samples and perturbation-based assays, structurally protecting four of the five datasets in the contamination audit (Norman, Replogle, Weinreb LARRY, sci-fate). Geneformer’s independent Genecorpus-30M additionally excluded immortalized and malignant cell lines; the documented inclusion and exclusion criteria for both Census and Genecorpus-30M are what make the Dim D contamination caveat assessable in the first place. Two contamination findings are noted for completeness: Arc State’s transition module was trained directly on Replogle K562 data [7, 24], which would preclude any future Arc State evaluation on Replogle under the current contamination policy; and the NeurIPS 2021 CITE-seq dataset (GSE194122) is confirmed in CELLxGENE Census, so its RNA component was likely seen by all Census-trained models, leaving Geneformer as the only contamination-free Dimension D control. VCBench flags these overlaps rather than excluding results.

### 3.2 Perturbation response prediction (Dimension A; Norman CRISPRa)

VCBench extends Ahlmann-Eltze et al. [1] on Dimension A in three ways: it adds Arc State under a corrected regime; it separates the 107 GEARS test perturbations into an additive-evaluable subset (71 perturbations with both single-gene components in training) and a novel subset (36 perturbations with at least one component held out); and it identifies the regime where foundation models add value the additive baseline cannot. On the additive-evaluable subset, the additive baseline achieved PRR 0.890, exceeding the best foundation model (Geneformer V2-316M, PRR 0.627) by 42% (Figure 2a; head-to-head: Geneformer 0.627, scGPT 0.545, Transcript-Former −0.165). Arc State, retrained under the correct GEARS split, produced PRR 0.402, falling below the mean baseline (0.579); training loss converged to 0.027 while validation loss diverged to approximately 0.40, indicating overfitting to 138 training perturbations. On the novel subset, where the additive baseline is structurally inapplicable, scGPT achieved PRR 0.420 (Figure 2b), demonstrating modest but non-trivial compositional generalization unavailable to the additive method. To our knowledge this regime has not been analyzed separately in prior benchmarks.

**Figure 2.**
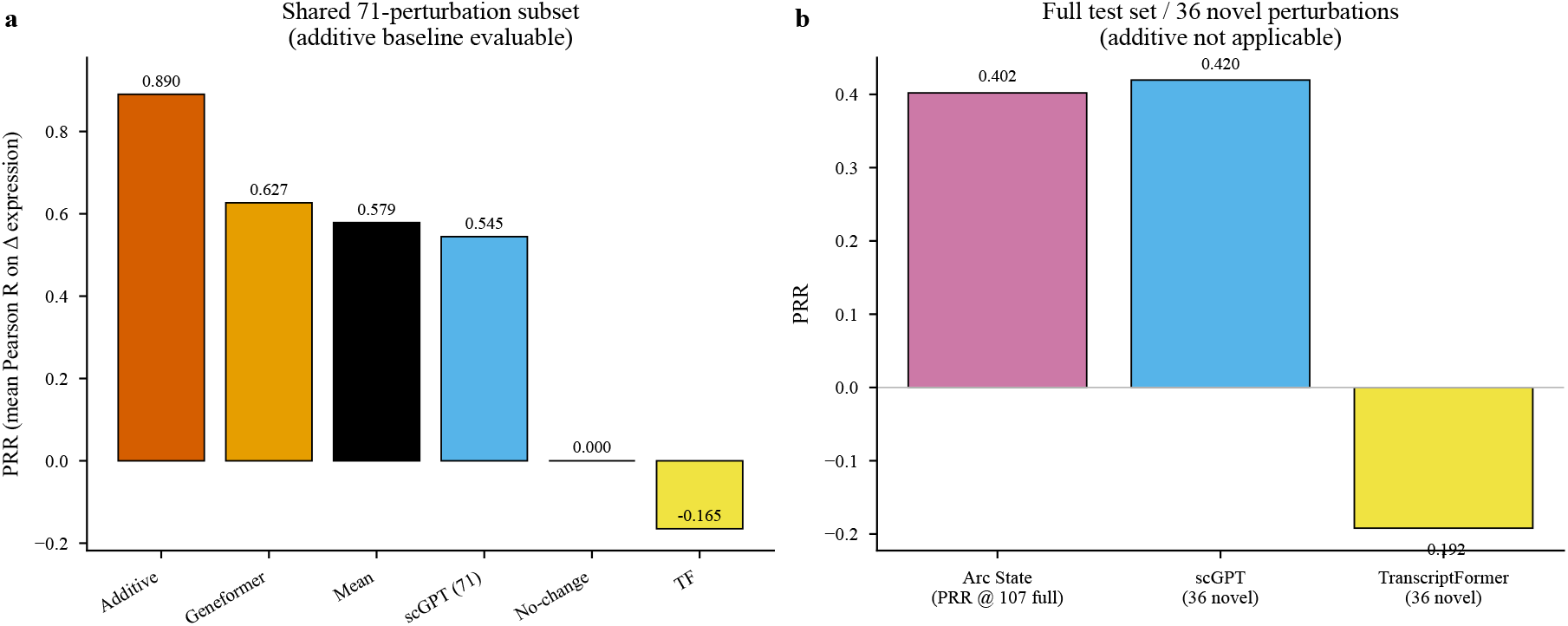
Perturbation prediction (Dimension A) results stratified by evaluability subset. (a) Shared 71-perturbation subset where the additive baseline is evaluable: additive baseline 0.890 substantially exceeds every foundation model (Geneformer 0.627, Mean 0.579, scGPT 0.545, no-change 0.000, TranscriptFormer −0.165). (b) Full test set / 36 novel perturbation subset where the additive baseline is structurally inapplicable: scGPT achieves PRR 0.420; TranscriptFormer achieves PRR −0.192; Arc State on the full 107-perturbation set is shown for reference (PRR 0.402).

To distinguish regime from representation in the cross-model comparison, we evaluated the same Geneformer V2-316M checkpoint in two matched regimes: the FT+D regime reported in Table 2 (PRR 0.627, *n* = 106; one perturbation excluded from fine-tuning under the GEARS split due to insufficient training cells) and a matched ZS+D regime using the pretrained model with tokendeletion perturbation and a ridge decoder (PRR 0.239, *n* = 107). The 0.39 PRR gap between regimes confirms that fine-tuning is a substantial contributor. However, Geneformer’s ZS+D result still substantially outperforms TranscriptFormer’s matched ZS+D regime (PRR −0.174), placing the within-regime upper bound well above Transcript-Former’s anti-correlated predictions and establishing that TranscriptFormer’s negative PRR reflects a genuine representational property of its zero-shot embeddings on this task rather than a regime artifact (Figure 3). The same encoding that benefits cross-modal RNA-to-protein prediction (Dimension D) appears to lack the perturbation-relevant geometry required for direction-of-effect prediction; this is consistent with TranscriptFormer’s multispecies autoregressive pretraining objective, which optimizes for sequence-level molecular similarity rather than perturbation-conditioned response.

**Figure 3.**
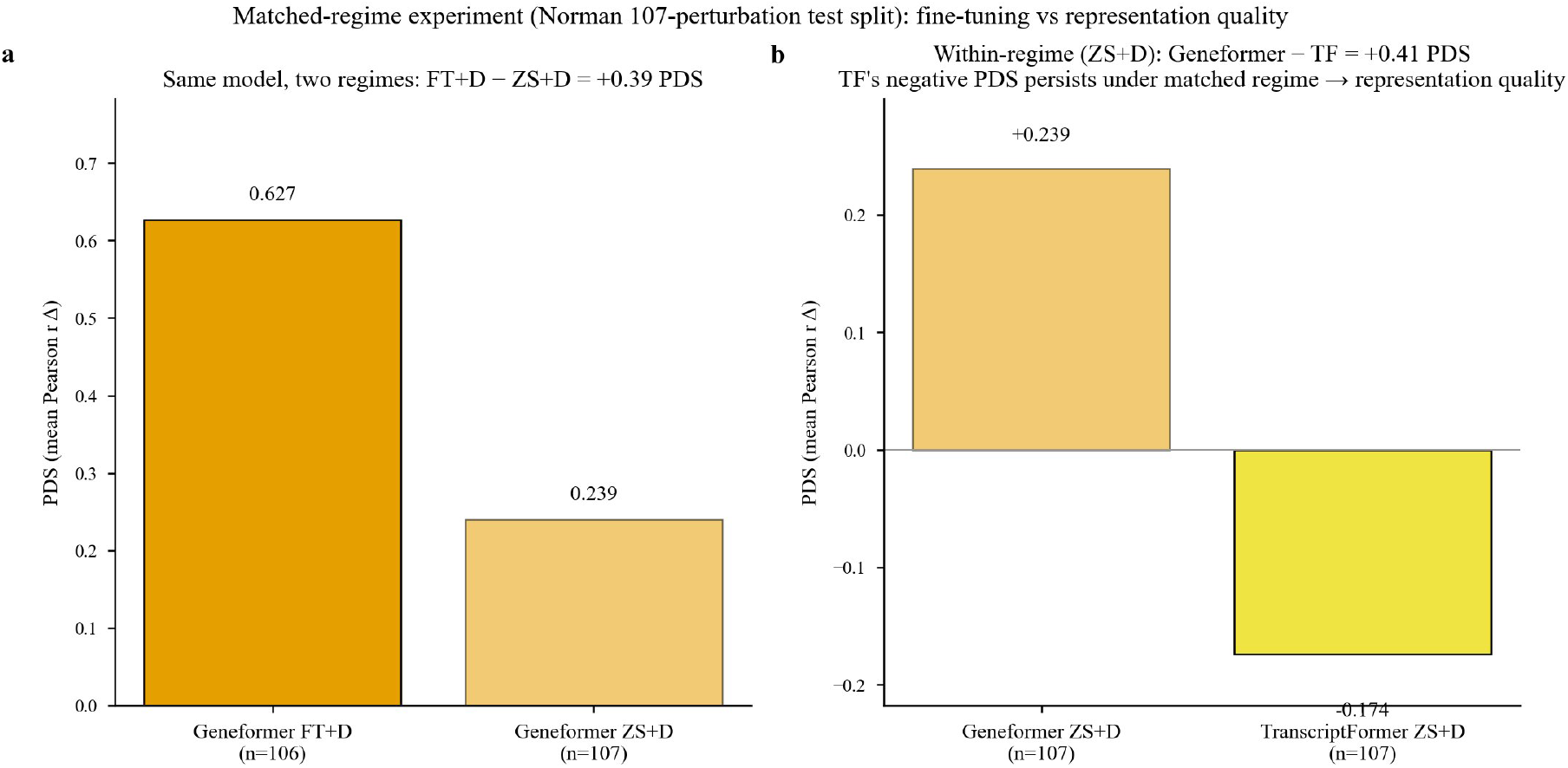
Matched-regime experiment on Norman 107-perturbation test split. (a) Geneformer evaluated under FT+D (*n* = 106) and ZS+D (*n* = 107) regimes; the 0.39 PRR gap quantifies the fine-tuning contribution. (b) Within the matched ZS+D regime, Geneformer (PRR 0.239) substantially exceeds TranscriptFormer (PRR −0.174), establishing that TranscriptFormer’s negative PRR reflects representation quality rather than a regime artifact.

### 3.3 Cross-species universality (Dimension B)

Cross-species cell-type transfer was evaluated under both a native-label protocol and a common-label-set protocol (see Methods). Under the native protocol, Geneformer’s aggregate macro F1 (0.181) appeared to exceed PCA+kNN’s (0.166), and TranscriptFormer’s lung+liver weighted F1 (0.347) appeared to exceed the matched baseline. Under the common-label-set protocol, both apparent leads disappear: PCA+kNN aggregate macro F1 is 0.497 (versus Geneformer 0.171); on lung+liver, PCA+kNN aggregate macro F1 is 0.456 and weighted F1 is 0.741, exceeding TranscriptFormer’s 0.351 and 0.560 on both metrics. Per-tissue under the common-label-set protocol, PCA+kNN exceeds every foundation model on lung, heart, kidney, and brain; on liver, TranscriptFormer (macro F1 0.495) marginally exceeds PCA+kNN (0.446), the only foundation-model-beats-baseline cell in the per-tissue matrix (Figure 4; Supplementary Table 2). The lung+liver aggregate (0.456 baseline vs 0.351 TF) thus averages over a tissue-level inversion. The native-protocol leads were artifacts of comparing macro F1 across models scored on differently-sized vocabularies; once matched, foundation-model performance falls below the linear baseline at the aggregate level on cross-species transfer, with the single liver exception noted above.

**Figure 4.**
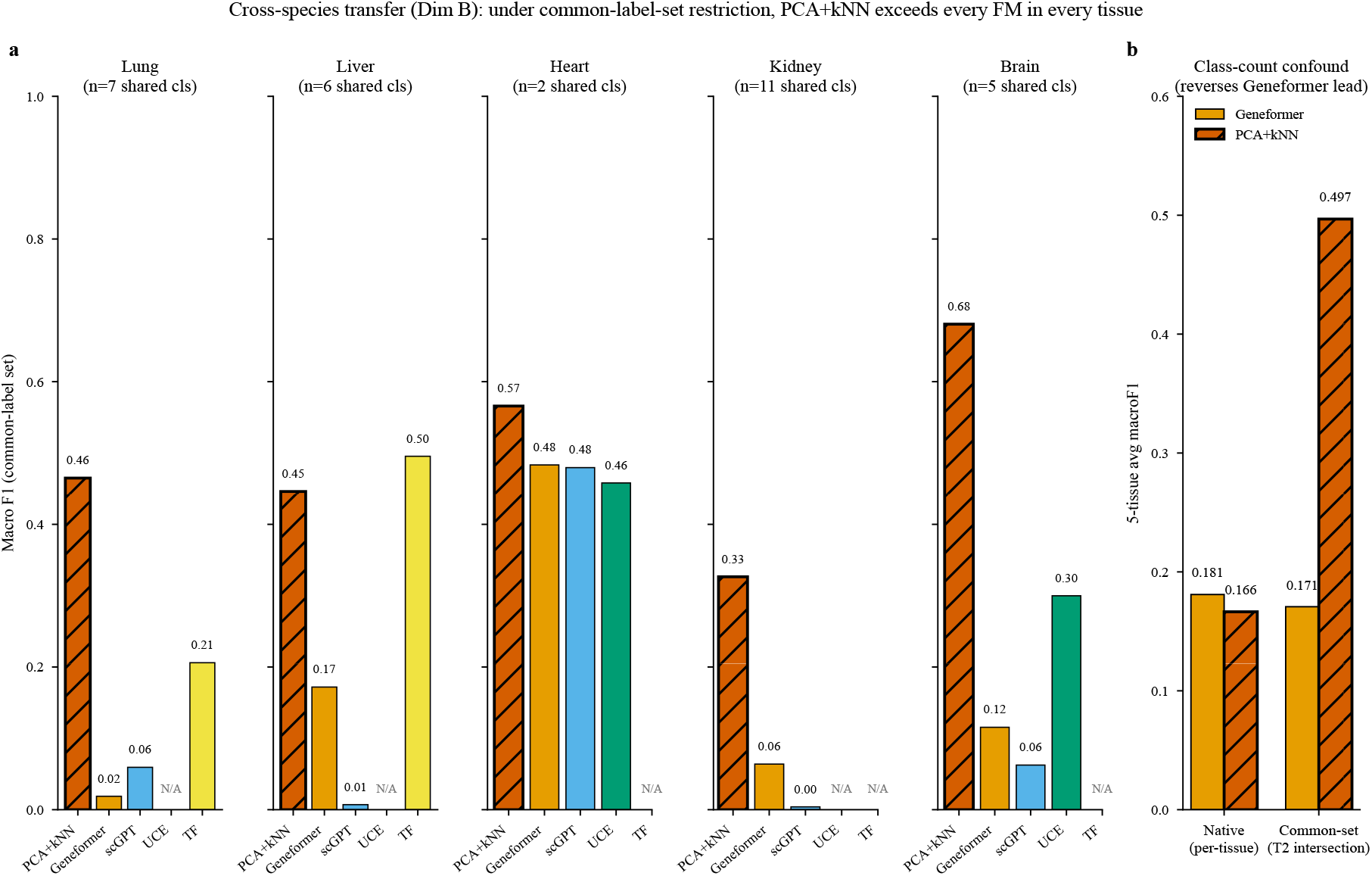
Cross-species transfer (Dimension B) under common-label-set restriction. (a) Per-tissue macro F1 for each foundation model and the PCA+kNN baseline restricted to the same cell-type vocabulary per tissue. PCA+kNN exceeds every foundation model on lung, heart, kidney, and brain; on liver, TranscriptFormer (0.495) marginally exceeds PCA+kNN (0.446). (b) Five-tissue aggregate comparison under native versus common-label-set protocol: the common-label-set protocol reverses the native-protocol Geneformer lead, revealing it as a class-count confound.

### 3.4 Gene regulatory network inference (Dimension C)

No foundation model passes the conjunctive AUPRC + EPR rule (Methods). Bootstrap-resampled 95% confidence intervals (1,000 paired iterations) on AUPRC place Geneformer (0.001) within the co-expression CI (0.004), with paired test BH *q* = 0.692; AUPRC overlap with the binding co-expression baseline cannot be rejected. scGPT’s AUPRC (0.003) is statistically distinguishable from co-expression (*q <* 0.001) but lies below it. On EPR, scGPT (20.05) clears every baseline (co-expression 15.50, pySCENIC 3.50, degree-null 1.13) by a wide margin but fails the conjunctive rule because of the AUPRC failure: scGPT’s gene-embedding geometry organizes high-confidence TF ⟶ target relationships better than chance even though the global precision-recall surface does not separate from co-expression. Geneformer additionally exhibits an AUROC/AUPRC dissociation (AUROC 0.626, AUPRC 0.001, EPR 0.000) consistent with attention capturing co-expression rather than unique regulatory signal: above-chance ranking on the full set coexists with zero true edges in the top-ranked predictions, replicating Kendiukhov [12] with different methodology and a different evaluation corpus (Figure 5). pySCENIC sits between the degree-null floor (AUPRC 0.0003) and co-expression (AUPRC 0.004), confirming that motif-pruned inference adds modest enrichment over random. TranscriptFormer is N/A on Dimension C (Methods); UCE and Arc State are N/A on architectural grounds.

**Figure 5.**
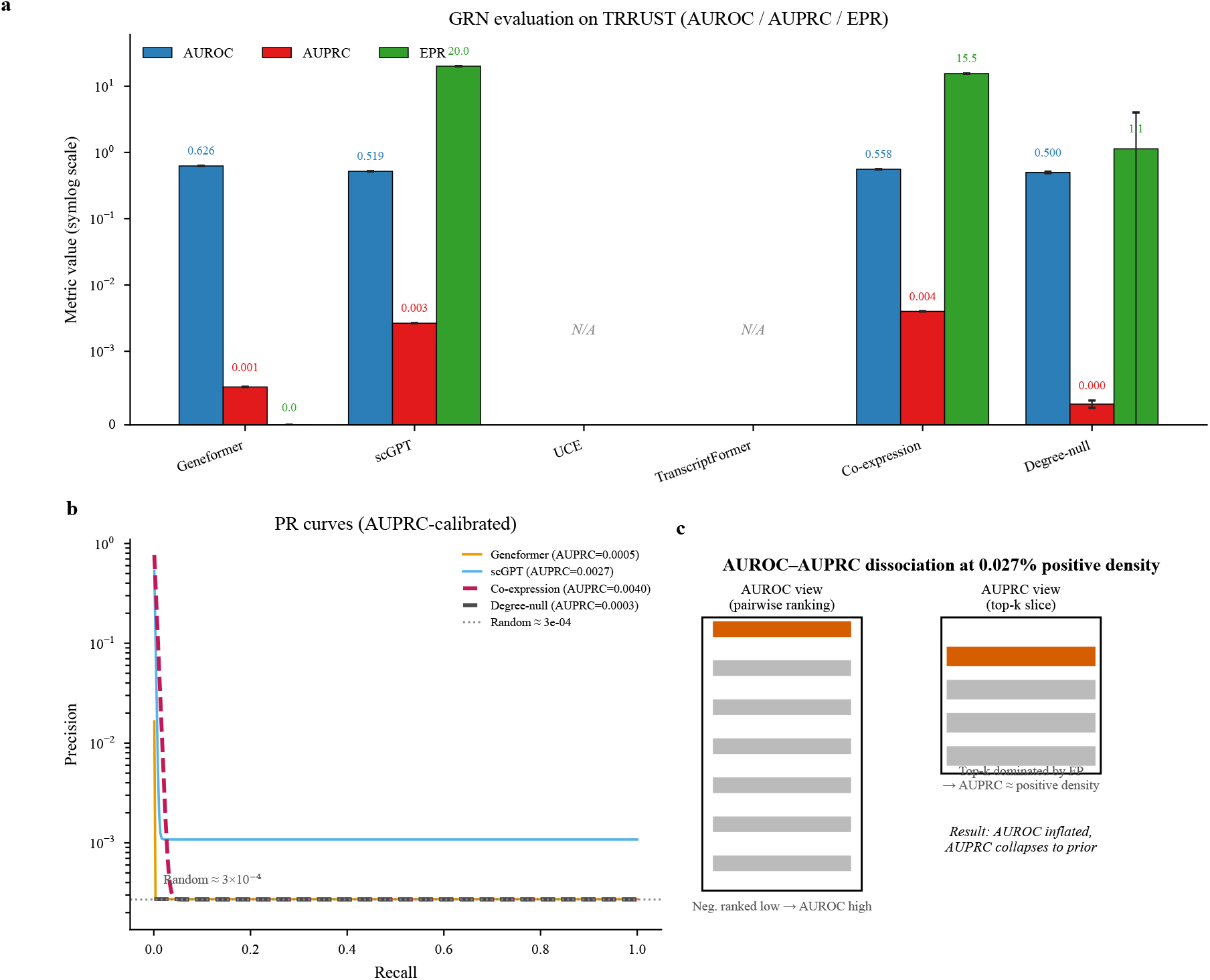
Gene regulatory network inference (Dimension C) on TRRUST. (a) Bar chart of AUROC, AUPRC, and EPR for each foundation model and baseline on symlog scale. (b) Precision-recall curves with AUPRC values per model; the random baseline at 3 × 10^*−*4^ reflects the positive-class density on TRRUST (0.027% true regulatory edges among evaluated pairs). (c) Mechanistic explanation of the AUROC-AUPRC dissociation pattern: above-chance pairwise ranking inflates AUROC while the top-*k* slice remains dominated by false positives, collapsing AUPRC to the prior.

### 3.5 Cross-modal prediction (Dimension D)

TranscriptFormer is the only foundation model to exceed every baseline, achieving per-protein Pearson 0.232 against the mean-celltype baseline’s 0.152, a 53% improvement. The other foundation models fall below the strongest baseline (UCE 0.132, scGPT 0.064, Geneformer 0.001, the latter near zero); Arc State lacks a public crossmodal API and is reported DNR. UCE in particular was designed for cross-species cell-type alignment rather than cross-modal prediction, and its Dimension D result is reported for capability-matrix completeness rather than as a comparison against its design targets. RNA-to-protein predictive sufficiency is therefore not a general property of foundation-model embeddings but depends on architectural choices, here mechanistically attributable to TranscriptFormer’s multi-species autoregressive pretraining, which appears to encode molecular-similarity features relevant to protein abundance. A contamination caveat applies: the NeurIPS 2021 CITE-seq dataset (GSE194122) is present in CELLxGENE Census and was likely seen by all four Census-trained models. Geneformer (Census-independent) achieving 0.001 on the same task indicates that exposure to the RNA component alone does not guarantee good performance, but does not rule out that Transcript-Former’s advantage is partly inflated by training-time exposure to GSE194122 expression patterns.

### 3.6 Temporal ordering (Dimension E)

The PCA+DPT baseline (Kendall’s *τ* -b 0.190, balanced accuracy 0.679) exceeds every foundation model. UCE approached but did not reach it (*τ* 0.136). TranscriptFormer produced weak positive signal with high variance (aggregate *τ* 0.046; sci-fate 0.051, Weinreb 0.041 ± 0.078 computed via 10 × 5,000-cell bootstrap after full-dataset diffusion-map eigensolution failed on Weinreb due to near-duplicate embeddings producing a degenerate graph Laplacian; Methods). Geneformer and scGPT produced negative or near-zero *τ* on average (− 0.017 and −0.057 respectively); on Weinreb specifically, scGPT achieved *τ* = − 0.103, indicating mild but consistent inversion of developmental time rather than failure to encode it. To our knowledge this is the first reproduction in an independent evaluation pipeline of the temporal-compression phenomenon (cells from different timepoints occupying overlapping embedding regions) described by Zhou et al. [25] in scGPT on Weinreb. The TranscriptFormer result is informative when considered jointly with its Dimension D advantage: the molecular-similarity encoding that benefits protein prediction also collapses temporally distinct cells onto near-identical embedding positions, destroying the spectral structure that diffusion pseudotime requires (Figure 6; Figure 7). To our knowledge, this Dimension D / Dimension E architectural tradeoff has not been characterized previously as a single-model dissociation.

**Figure 6.**
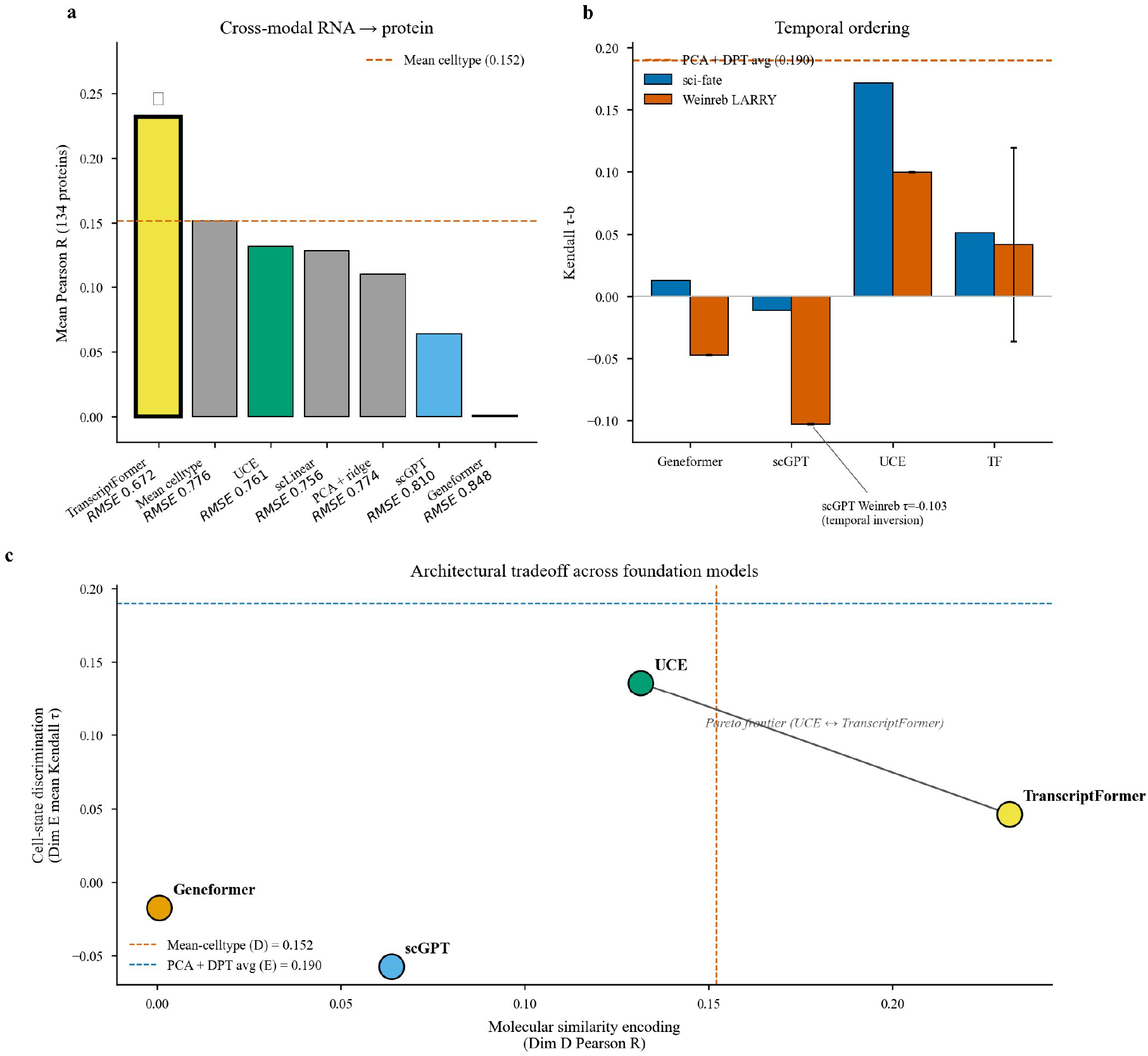
Cross-modal prediction, temporal ordering, and the architectural tradeoff between them. (a) Cross-modal RNA ⟶ protein per-protein Pearson R (134 ADTs) on NeurIPS 2021 CITE-seq. TranscriptFormer (⋆) is the only foundation model to exceed the mean-celltype baseline (0.152). RMSE values annotated for each entry. (b) Temporal ordering Kendall *τ* -b on sci-fate and Weinreb LARRY. PCA+DPT (dashed line, average 0.190) exceeds every foundation model. scGPT on Weinreb shows *τ* = −0.103, indicating systematic temporal inversion. (c) Architectural tradeoff scatter showing each foundation model’s Dimension D Pearson R against its Dimension E mean Kendall *τ*. UCE and TranscriptFormer occupy a Pareto frontier in opposite directions; the molecular-similarity encoding that makes TranscriptFormer succeed on Dimension D is the same property that produces its near-zero Dimension E result via spectral collapse (Figure 7).

### 3.7 Composite capability assessment

Under the pre-registered VC Level mapping, all five foundation models achieve Level 1: each exceeds the trivial baseline on at least one scored dimension (per the per-dimension scores in Table 2). One model achieves Level 2: TranscriptFormer on cross-modal prediction. No model achieves Level 3 (Figure 8). The pattern is consistent: linear and nearest-neighbor baselines match or exceed every foundation model on every dimension where simple baselines apply, with one architecturally-localized exception on Dimension D. Joint inspection of per-dimension scores surfaces a second cross-dimensional pattern beyond the TranscriptFormer Dimension D / Dimension E tradeoff: scGPT shows a fine-scale-versus-population-scale dissociation, with the highest EPR of any evaluated method on Dimension C (20.05) coexisting with one of the lowest per-protein Pearson correlations on Dimension D (0.064). The two findings are consistent with a model that learns local gene-gene relationships well and global cell-state geometry poorly. We treat this observation as illustrative rather than systematic; a single second example does not establish that the framework reliably uncovers such patterns, and we expect their density to grow as more models are added.

**Figure 7.**
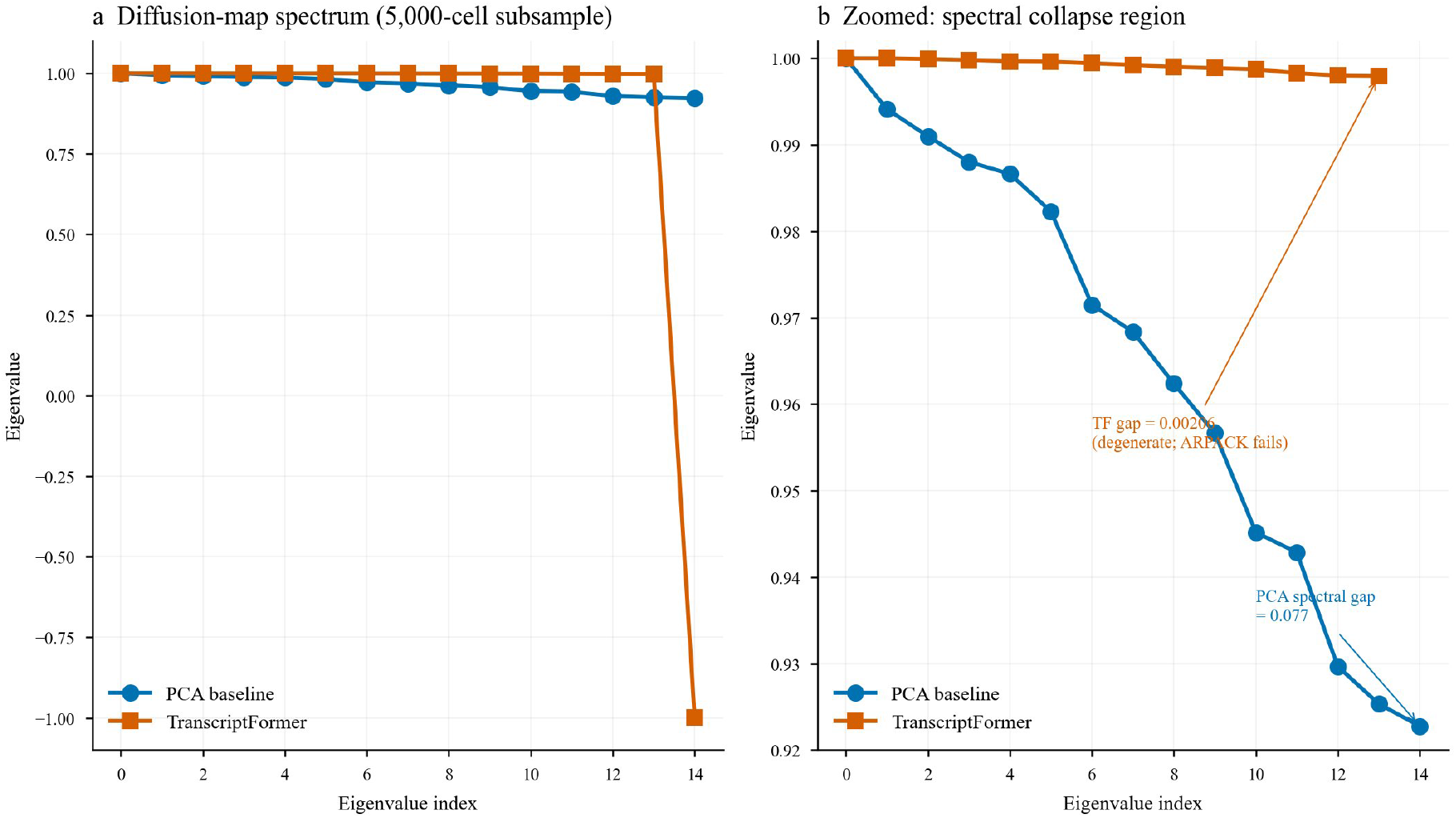
Spectral analysis of TranscriptFormer embeddings on Weinreb. (a) Diffusion-map graph Laplacian eigenvalue spectrum (5,000-cell subsample) for TranscriptFormer (orange) versus a PCA baseline (blue). (b) Zoomed view of the spectral collapse region: the PCA baseline exhibits a healthy spectral gap of 0.077; TranscriptFormer’s gap is 0.00206, an effectively degenerate spectrum that prevents ARPACK convergence.

**Figure 8.**
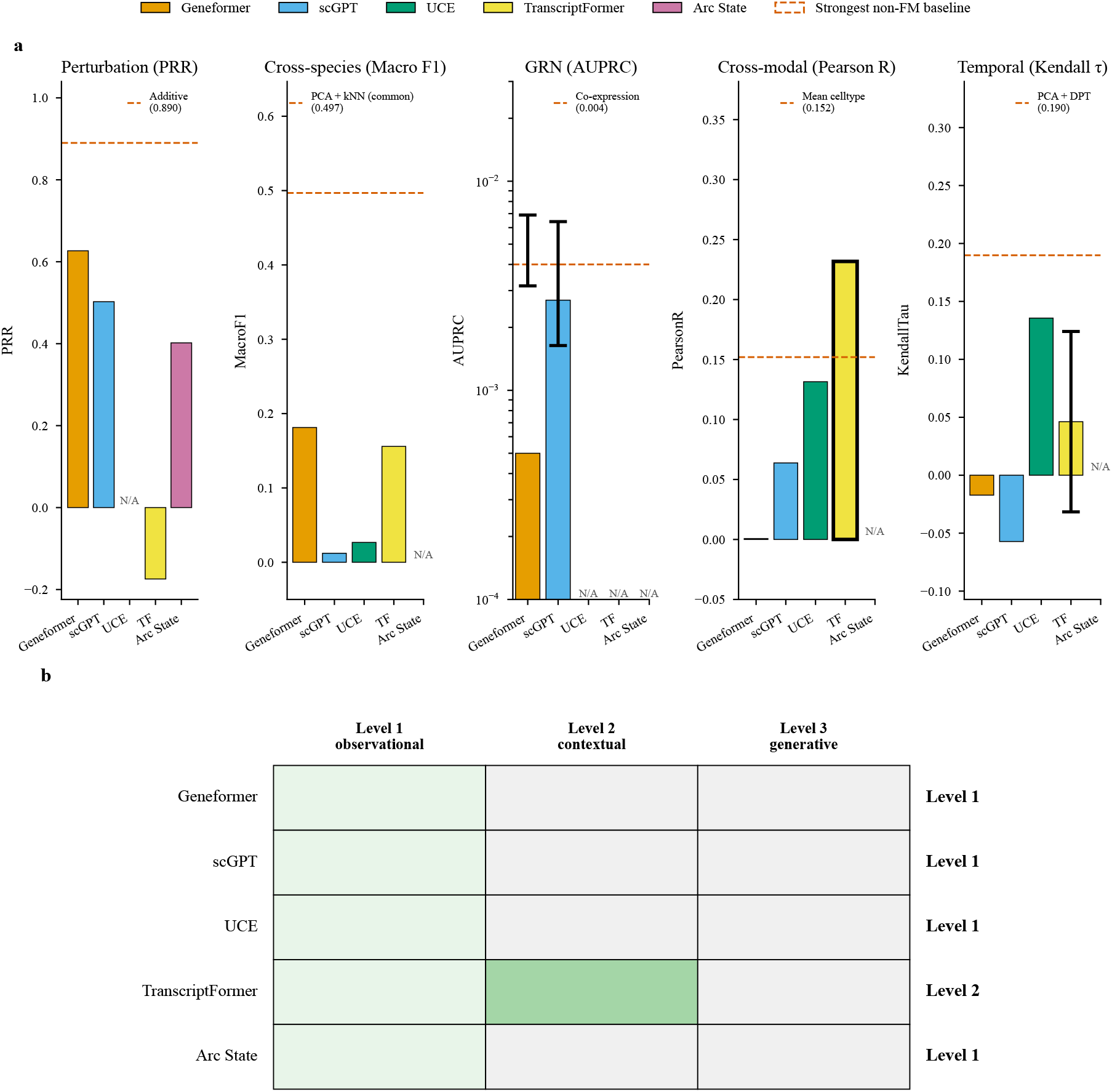
VC Level assignments across five foundation models and five evaluation dimensions. (a) Per-dimension score comparison on the primary dataset for each dimension (Norman for Dimension A; matched-tissue average under common-label-set protocol for Dimension B; BEELINE/TRRUST for Dimension C; NeurIPS CITE-seq for Dimension D; sci-fate and Weinreb LARRY average for Dimension E). For each dimension the bar chart shows the primary metric value for each foundation model alongside the strongest non-foundation-model baseline (dashed reference line). The bold-outlined bar with star marks the only Level-2 cell. Error bars on Dimension C show 95% confidence intervals from 1,000 paired bootstrap iterations on AUPRC; error bars on TranscriptFormer Weinreb show bootstrap standard deviation across 10 random 5,000-cell subsamples. Other entries are point estimates from a single evaluation. N/A entries indicate models architecturally incompatible with the dimension. (b) VC Level assignment grid. All five foundation models reach Level 1; one (TranscriptFormer) reaches Level 2 on cross-modal prediction; none reaches Level 3.

### 3.8 Two dimensions remain structurally untestable as end-to-end tasks

Multi-scale integration requires propagating predictions across DNA sequence, gene expression, cellular phenotype, and tissue architecture within a single architecture. No current model approaches this. Enformer [41] and Borzoi [42] predict expression from DNA without modeling cell state; Nicheformer [43] models spatial niches without molecular sequence. Thornburg et al. [26], extending earlier whole-cell simulation work [27] on Mycoplasma genitalium, demonstrated what genuine multi-scale integration requires: a single model tracking ribosome assembly, metabolic flux, DNA replication, and cell division simultaneously, with reaction stoichiometry enforcing mass balance, requiring six days on two dedicated GPUs for a 493gene minimal cell. No paired dataset spans DNA through tissue for the same biological system at single-cell resolution.

In silico experimentation has two components with distinct testability status. The closed-loop labin-the-loop component is structurally untestable in a static benchmark by definition. The epistemic-calibration prerequisite (a model that reports when it does not know) is testable, and VCBench tests it via a spread-error correlation probe: for each model’s perturbation predictions on Norman, the Spearman rank correlation between per-perturbation LFC spread and per-perturbation MAE is computed. Neither evaluated model shows a detectable correlation (Geneformer *ρ* = −0.119, *p* = 0.225; scGPT *ρ* = +0.131, *p* = 0.177; *n* = 106-107; Supplementary Table 8). The LFC spread of either model’s point prediction carries no information about where the prediction will err. This is necessary but not sufficient for epistemic calibration: a model with real predictive-interval calibration would pass this test, but passing it would not establish real uncertainty. A direct test would require distributional output heads, ensembles, or conformal wrappers, which we recommend for future benchmark versions.

## 4 Discussion

VCBench operationalizes five of seven virtual cell capability dimensions and evaluates five foundation models against this framework. The results expose a consistent pattern: simple baselines match or exceed the best foundation model on four of five dimensions (perturbation prediction, cross-species transfer under matched class sets, GRN inference, temporal ordering), one foundation model shows a genuine but narrow architectural advantage (TranscriptFormer on cross-modal prediction), and no model achieves Level 3. Where models can be scored, simple baselines remain competitive; where they cannot be scored, the gap is architectural rather than a matter of scale.

Joint inspection across dimensions surfaces patterns invisible to single-task benchmarks. Beyond TranscriptFormer’s Dimension D advantage and Di-mension E spectral collapse (Figure 6), two further dissociations are worth flagging. Geneformer on Dimension C produces above-chance edge ranking (AUROC 0.626) coexisting with near-zero precision in the top-ranked predictions (AUPRC 0.001, EPR 0.000), replicating the Kendiukhov [12] finding that attention captures co-expression rather than unique regulatory signal; an attention-derived GRN can rank true edges above random while still failing to concentrate them at the top, which is the regime where downstream uses of an inferred network actually live. scGPT exhibits a complementary pattern: its EPR on Dimension C (20.05) is the highest of any evaluated method while its per-protein Pearson on Dimension D (0.064) is among the lowest, consistent with a model that learns local gene-gene relationships well and global cell-state geometry poorly. We treat these joint observations as illustrative rather than systematic; a small number of cross-dimensional dissociations does not establish that the framework reliably uncovers them, and we expect their density to grow as more models are added. The finding worth carrying forward is methodological: cross-dimensional dissociations are detectable only because dimensions are scored with shared protocols and reported jointly.

A natural response to this pattern is that foundation models were designed for representation learning, not interventional prediction [1]. This objection has merit for any benchmark that evaluates models outside their intended scope. However, the virtual cell frameworks that motivate these models explicitly demand interventional capability: Bunne et al. [2] described Manipulator Virtual Instruments that predict cell state transitions, Noutahi et al. [9] placed perturbation prediction as the first of three core capabilities, and the Arc Institute’s Virtual Cell Challenge [8] was framed as “a Turing test for the virtual cell.” VCBench holds models to the standard their proponents have set. If that standard is premature, the gap itself is a finding worth documenting.

That said, VCBench evaluates a specific and demanding slice of virtual cell capability. The same models that fail perturbation prediction or temporal ordering may perform well on cell-type annotation, batch correction, or data integration, tasks that are also part of the virtual cell vision but are not tested here because they have been benchmarked extensively elsewhere [11, 19, 37]. The claim that “the gap is structural” applies to the interventional and cross-dimensional capabilities VCBench measures, not to foundation model utility in general.

A related objection concerns metric choice. Viñas Torné et al. [28] showed that commonly used perturbation metrics can be inflated by systematic variation shared across perturbations. PRR is susceptible to this inflation: the mean-prediction baseline achieves PRR 0.579 on the additive-evaluable Norman subset, well above zero, illustrating exactly the systematic-variation floor Viñas Torné describes. VCBench mitigates this by reporting all foundation-model PRR values alongside this mean baseline, so that the comparison the reader sees is foundation-model-versus-baseline rather than against zero. Future versions should incorporate the rank-based decomposition of Viñas Torné and the PerturBench framework [29] to further separate perturbation-specific prediction from dataset-level artifacts.

VCBench evaluates each model using the strongest protocol available to it (fine-tuning where supported on Dimension A; zero-shot embedding probes on Dimensions B, D, E; internal-state extraction on Dimension C) rather than forcing a uniform regime. The capability matrix annotates each entry with its evaluation regime. Where models share a regime within a dimension (cross-model comparisons), regimevs-representation ambiguity is addressed directly by the matched-regime experiment on Dimension A (Figure 3): TranscriptFormer’s negative PRR is established as a representation-quality finding rather than a regime artifact.

Dimensions D and E are proxy tasks rather than direct measurements of the underlying capabilities. A linear probe on RNA embeddings can establish predictive sufficiency for protein abundance (Dimension D) but not representational invariance in the sense of Bunne et al. [2]; diffusion pseudo-time can fail on embeddings whose temporal information is present but spectrally inaccessible (the TranscriptFormer Dimension E result is a concrete instance of this). Both proxies are conservative tests that can demonstrate failure but may underestimate capability.

VCBench has three principal limitations that scope its claims. First, all scored dimensions operate on transcriptomic data; classical estimates put the mRNA-explained fraction of protein-level variance at ~40% in mammalian cells [30], with noise-corrected analyses revising this to 56–81% in mammals [31] and *>*85% in yeast [32], though substantial post-transcriptional contribution remains. Virtual cells must eventually span proteomics, metabolomics, chromatin state, and spatial organization that lie outside the present benchmark. Second, models are evaluated in publicly released configurations; emerging architectures and alternative training regimes may shift the empirical landscape. Third, two of seven dimensions remain structurally untestable. Operational caveats on the contamination audit and spread-error probe coverage are documented in Supplementary Note 2.

The five scored dimensions implicitly specify architectural requirements for a next-generation virtual cell model: a multi-species tokenizer extended beyond protein-coding genes, explicit temporal conditioning or trajectory-aware pretraining objectives, paired multi-omic pretraining corpora, calibrated uncertainty for active-learning loops, and explicit regulatory training objectives in place of post-hoc attention extraction [12]. These requirements are individually achievable but have not been combined in a single architecture. The spread-error correlation probe provides empirical grounding for the calibration requirement: the absence of even minimal spread-error correlation in both tested models is a measured deficit, not a theoretical concern. Without calibrated uncertainty, the active-learning loops central to the virtual cell vision [2, 9] cannot be closed regardless of improvements in raw predictive accuracy.

The VC Level assignments depend on threshold choices adopted from Noutahi et al. [9]; these are deliberately lenient (Level 2 requires exceeding the strongest baseline on only one dimension). If Level 2 required exceeding on two dimensions, no model would qualify; if Level 3 required four dimensions, no model would be within reach. The Level framework should be tightened as the field matures.

The contamination audit revealed a systemic transparency deficit: no evaluated model publishes a complete cell-level training manifest. We propose the VCBench Contamination Reporting Schema v1 (Supplementary Note 1) as a minimal standard: MD5 hashes of pretraining cell barcodes; a standardized `anndata.obs` flag marking cells as training, validation, or held-out; and accession-level manifests indicating which datasets were included. These are individually low-effort and would together permit exact contamination detection rather than the structural-asymmetry audit currently required. Natural language processing (NLP) addressed the analogous problem with ngram-overlap detection and canonical decontamination protocols [35, 36]; single-cell biology requires an analogous standard. Arc Institute’s published training-data documentation [7], which enumerates the constituent datasets of the Arc State transition module training corpus, provides the closest current implementation of the proposed schema and a natural starting point for community adoption.

Beyond the empirical findings, this work contributes three methodological tools that are reusable independently of the specific models evaluated here. The common-label-set protocol (Eq. 4) corrects a class-count confound that inflates apparent foundation-model performance on cross-species transfer under heterogeneous tokenizer vocabularies; it applies to any benchmark where evaluated methods admit different class sets. The spread-error correlation probe (Eq. 10) is a necessary-but-not-sufficient calibration test that detects models lacking even minimal coupling between predicted output spread and prediction error; it applies to any regression-style perturbation predictor. The Contamination Reporting Schema is independent of the benchmark itself and could be adopted as a community-level transparency standard. We release all three alongside the benchmark code so that they can be adopted, extended, or contested separately from the headline results.

VCBench should evolve as the field advances, following the GLUE-to-SuperGLUE precedent [33, 34]. As multi-modal architectures mature, the cross-modal dimension should transition from proxy to direct evaluation; the temporal dimension should expand beyond ordering to time-indexed prediction using metabolic-labeling datasets; and the two structurally untestable dimensions should be revisited as hybrid architectures and prospective validation protocols emerge. The finding that current models score on a minority of seven dimensions does not diminish the virtual cell vision; it calibrates expectations, identifies specific capability gaps, and provides measurement infrastructure to track progress.

## 5 Data and code availability

### Code availability

All analysis code is available at github.com/AppliedScientific/VCBench (release tag v1.0.0, commit 079355d, archived at github.com/AppliedScientific/VCBench/releases/tag/v1.0.0) under an MIT license. The VCBench Contamination Reporting Schema v1 is released within the same repository.

### Model availability

The corrected Arc State Norman fine-tuned checkpoint, both runs’ raw evaluation AnnData files, and the superseded checkpoint from our earlier (leaked-configuration) run with its training metrics are publicly available at huggingface.co/VibeCodingScientist/arc-state-norman-gears-corrected(pinned revision `34d20125`). Paste-able reproduction snippets that recover the canonical PRR = 0.402 and the inflated PRR = 0.949 in under 5 minutes on CPU are in `forensic_artifacts/README.md` of that repository. The fine-tuned scGPT and Geneformer V2-316M checkpoints are released at the same HuggingFace organization.

### Data availability

Datasets used for evaluation are publicly available at the GEO accessions documented in Methods (Norman: GSE133344; CELLxGENE Census: cellxgene.cziscience.com; NeurIPS 2021 CITE-seq: GSE194122; Weinreb LARRY: GSE140802; sci-fate: GSE131351). Pre-computed model embeddings and predictions for every (model, dataset) cell in the capability matrix are released at the same HuggingFace organization to enable rapid reproduction without re-running expensive forward passes.

## Supplementary Information

**Supplementary Table 1.** Wall-clock evaluation time per model and dimension. All times on a single NVIDIA A100 80 GB GPU. Times include embedding extraction and evaluation but exclude one-time pretrained weight downloads. Format: H:MM:SS. Baseline computations (PCA+kNN, PCA+ridge, PCA+DPT, co-expression, additive, mean-prediction, scLinear, mean-celltype) completed in under 10 minutes total across all dimensions on CPU.

| Model | Dimension A | Dimension B | Dimension C | Dimension D | Dimension E |
| --- | --- | --- | --- | --- | --- |
| Geneformer V2-316M | 1:00:00* | 0:00:16 | 0:03:21 | 0:02:00 | 0:00:28 |
| scGPT (fine-tuned) | 0:00:04‡ | 0:02:45 | 0:00:15 | 0:49:34 | 0:00:01‡ |
| UCE 33-layer | N/A | 10:18:00 | N/A | 1:58:00 | 1:12:00 |
| TranscriptFormer | 3:26:00 | 9:48:00§ | 0:05:00 | 0:01:34 | 0:00:21 |
| Arc State | 7:22:00¶ | N/A | N/A | DNR | DNR |

\*Wall-clock estimated from log cadence; full re-run not attempted given resource constraints. ‡Cached short-circuit run; original training time folded into the parent master pipeline. §Sum of four attempts including two OOM retries (TranscriptFormer Dimension B completed on lung + liver only). ¶Arc State 110M training on the corrected GEARS split (`max_steps = 40,000` reached).

**Supplementary Table 2.** Per-tissue cross-species results. Breakdown by tissue for all models on Dimension B, enabling matched-tissue comparisons beyond the aggregate numbers reported in Table 2. All tissues use 50,000 human + 50,000 mouse cells sampled from CELLxGENE Census and the 15,847 one-to-one high-confidence orthologs from the joint Ensembl mapping. The `n_shared_celltypes` column reports the intersection of ontology labels observed after each model’s own tokenizer/filter is applied. Geneformer drops cells whose in-vocabulary gene count falls below its rank-based filter, which reduces the label intersection relative to scGPT, UCE, and TranscriptFormer that operate on the full 50K-per-tissue sample. This reflects a genuine preprocessing difference, not an export error. Arc State is structurally N/A on Dimension B: its tokenizer lacks the ortholog mapping required for cross-species transfer. UCE was evaluated on heart and brain only; TranscriptFormer was evaluated on lung and liver only, reflecting species-vocabulary compatibility and compute budget constraints documented in Methods.

| Model | Tissue | Macro F1 | Weighted F1 | n_shared_celltypes |
| --- | --- | --- | --- | --- |
| Geneformer V2-316M | Lung | 0.0188 | 0.0093 | 7 |
| Geneformer V2-316M | Liver | 0.1668 | 0.2383 | 6 |
| Geneformer V2-316M | Heart | 0.5645 | 0.9109 | 2 |
| Geneformer V2-316M | Kidney | 0.0679 | 0.3790 | 11 |
| Geneformer V2-316M | Brain | 0.0886 | 0.0524 | 5 |
| scGPT (fine-tuned) | Lung | 0.0001 | 0.0000 | 54 |
| scGPT (fine-tuned) | Liver | 0.0431 | 0.4240 | 17 |
| scGPT (fine-tuned) | Heart | 0.0075 | 0.0212 | 26 |
| scGPT (fine-tuned) | Kidney | 0.0085 | 0.0864 | 44 |
| scGPT (fine-tuned) | Brain | 0.0013 | 0.0041 | 69 |
| UCE 33-layer | Lung | N/A | N/A | N/A |
| UCE 33-layer | Liver | N/A | N/A | N/A |
| UCE 33-layer | Heart | 0.0299 | 0.0858 | 26 |
| UCE 33-layer | Kidney | N/A | N/A | N/A |
| UCE 33-layer | Brain | 0.0235 | 0.0666 | 69 |
| TranscriptFormer | Lung | 0.0849 | 0.0867 | 54 |
| TranscriptFormer | Liver | 0.2269 | 0.6071 | 17 |
| TranscriptFormer | Heart | N/A | N/A | N/A |
| TranscriptFormer | Kidney | N/A | N/A | N/A |
| TranscriptFormer | Brain | N/A | N/A | N/A |
| Arc State | all tissues | N/A | N/A | N/A |

**Supplementary Table 3.** Per-dataset temporal ordering results. Breakdown by dataset (sci-fate vs Weinreb LARRY) for all models on Dimension E, showing Kendall’s *τ* -b and kNN balanced accuracy separately for each dataset rather than only the average reported in Table 2. The *τ* std (boot) column is non-empty only when Kendall *τ* was computed over bootstrap replicates rather than on the full matrix; single-shot runs report Kendall *p* instead. ^*†*^ TranscriptFormer Weinreb: the full 49,008-cell ARPACK eigendecomposition did not converge; values are mean *±*std over 10 random subsamples of 5,000 cells, and Kendall *p* is therefore not available. Arc State is N/A on Weinreb (tokenizer lacks the mouse-gene mapping required for LARRY) and DNR on sci-fate (not executed in this evaluation).

| Model | Dataset | Kendall $\tau$ | $\tau$ std | Kendall $p$ | kNN bal. acc. | $n_{\text{cells}}$ | $n_t$ |
| --- | --- | --- | --- | --- | --- | --- | --- |
| Geneformer V2-316M | sci-fate | 0.0128 | — | $1.49 \times 10^{-1}$ | 0.4094 | 6,567 | 4 |
| Geneformer V2-316M | Weinreb | -0.0473 | — | $1.45 \times 10^{-41}$ | 0.6977 | 49,008 | 3 |
| scGPT (fine-tuned) | sci-fate | -0.0115 | — | $1.96 \times 10^{-1}$ | 0.4386 | 6,567 | 4 |
| scGPT (fine-tuned) | Weinreb | -0.1028 | — | $9.30 \times 10^{-190}$ | 0.6668 | 49,008 | 3 |
| UCE 33-layer | sci-fate | 0.1715 | — | $6.79 \times 10^{-83}$ | 0.6143 | 6,567 | 4 |
| UCE 33-layer | Weinreb | 0.0997 | — | $1.73 \times 10^{-178}$ | 0.7198 | 49,008 | 3 |
| TranscriptFormer | sci-fate | 0.0510 | — | $9.98 \times 10^{-9}$ | 0.5495 | 6,567 | 4 |
| TranscriptFormer | Weinreb<sup>†</sup> | 0.0414 | 0.0779 | N/A | 0.3506 | 5,000 | 3 |
| PCA + DPT | sci-fate | 0.2269 | — | $1.55 \times 10^{-143}$ | 0.6199 | 6,567 | 4 |
| PCA + DPT | Weinreb | 0.1527 | — | $< 10^{-300}$ | 0.7375 | 49,008 | 3 |
| Arc State | sci-fate | DNR | — | DNR | DNR | — | — |
| Arc State | Weinreb | N/A | — | N/A | N/A | — | — |

**Supplementary Table 4.** Per-cell contamination rationale. Extended rationale for the classifications summarized in Supplementary Table 5. Arc State is decomposed into its Embedding (SE) and State Transition (ST) modules, which have different training corpora and therefore different contamination profiles. Risk categories: *Confirmed*, dataset is documented in the model’s training corpus; *Likely*, dataset meets the inclusion criteria of the model’s training platform with no exclusion documented; *Unlikely*, dataset fails documented exclusion criteria; *N/A*, the model module is not applicable to the evaluation dimension for that dataset. Risk assessments are based on published training data documentation (paper supplementary materials, data repository manifests) and platform-level schema (CELLxGENE Census inclusion/exclusion criteria). Where training manifests are incomplete (scGPT, UCE, TranscriptFormer), risk is assessed from platform-level inclusion criteria rather than cell-level overlap detection, which is currently infeasible given the absence of published cell-level training manifests from any evaluated foundation model. Datasets: Norman GSE133344; Replogle Figshare+ `10.25452/figshare.plus.20029387;` NeurIPS CITE-seq GSE194122; Weinreb LARRY GSE140802; sci-fate GSE131351.

| Model | Dataset | Risk | Rationale |
| --- | --- | --- | --- |
| Geneformer | Norman | Unlikely | Genecorpus-30M excludes cell lines; Norman uses K562 cell line |
| Geneformer | Replogle | Unlikely | Genecorpus-30M excludes cell lines; Replogle uses K562/RPE1 |
| Geneformer | CITE-seq | Unlikely | Independent corpus (pre-Census); bone marrow may be included but not ADT data |
| Geneformer | Weinreb | Unlikely | Genecorpus-30M is human-only; Weinreb is mouse HSPCs |
| Geneformer | sci-fate | Unlikely | A549 cell line excluded from Genecorpus |
| scGPT | Norman | Unlikely | Census excludes cell culture and perturbation assays |
| scGPT | Replogle | Unlikely | Census excludes cell culture and perturbation assays |
| scGPT | CITE-seq | Likely | GSE194122 present in Census 2023-05-15; RNA portion may overlap |
| scGPT | Weinreb | Unlikely | In vitro culture excluded from Census inclusion criteria |
| scGPT | sci-fate | Unlikely | A549 cell line excluded from Census |
| UCE | Norman | Unlikely | Census excludes cell culture |
| UCE | Replogle | Unlikely | Census excludes cell culture |
| UCE | CITE-seq | Likely | Census-derived training data; GSE194122 likely included |
| UCE | Weinreb | Unlikely | In vitro culture excluded |
| UCE | sci-fate | Unlikely | Cell line excluded |
| TranscriptFormer | Norman | Unlikely | Virtual Cells corpus excludes cell culture |
| TranscriptFormer | Replogle | Unlikely | Virtual Cells corpus excludes perturbation assays |
| TranscriptFormer | CITE-seq | Likely | CZI builds both Census and TranscriptFormer; high overlap likelihood |
| TranscriptFormer | Weinreb | Unlikely | In vitro culture excluded |
| TranscriptFormer | sci-fate | Unlikely | Cell line excluded |
| Arc State SE | Norman | Unlikely | Census excludes cell culture |
| Arc State SE | Replogle | Unlikely | Census excludes cell culture |
| Arc State SE | CITE-seq | Likely | Census Jan 2025 likely includes GSE194122 |
| Arc State SE | Weinreb | Unlikely | In vitro culture excluded |
| Arc State SE | sci-fate | Unlikely | Cell line excluded |
| Arc State ST | Norman | Unlikely | Not listed in Arc State ST training data |
| Arc State ST | Replogle | **Confirmed** | Replogle K562 explicitly in Arc State ST training data list |
| Arc State ST | CITE-seq | N/A | ST model does not evaluate on Dimension D (cross-modal) |
| Arc State ST | Weinreb | N/A | ST model does not evaluate on Dimension E (temporal) |
| Arc State ST | sci-fate | N/A | ST model does not evaluate on Dimension E (temporal) |

**Supplementary Table 5.** Contamination audit matrix. Contamination audit matrix for five foundation models against VCBench evaluation datasets. Risk levels: *Confirmed*, dataset documented in training corpus; *Likely*, dataset meets inclusion criteria of training data platform with no exclusion documented; *Unlikely*, dataset fails documented exclusion criteria; *Unknown*, insufficient training manifest. CELLxGENE Census excludes cell-culture tissue type and perturbation-based assays, providing structural protection for four of five evaluation datasets against models trained on Census. Arc State is split into its Embedding (SE) and State Transition (ST) modules, which have different training corpora. Per-cell rationale for each risk classification is provided in Supplementary Table 4.

| Model | Norman | Replogle | NeurIPS CITE-seq | Weinreb LARRY | sci-fate |
| --- | --- | --- | --- | --- | --- |
| Geneformer V2-316M | Unlikely | Unlikely | Unlikely | Unlikely (mouse) | Unlikely |
| scGPT | Unlikely (Census excl.) | Unlikely (Census excl.) | Likely (in Census 2023-05-15) | Unlikely (Census excl.) | Unlikely (Census excl.) |
| UCE 33-layer | Unlikely | Unlikely | Likely (Census-derived) | Unlikely | Unlikely |
| TranscriptFormer | Unlikely | Unlikely | Likely (CZI builds both) | Unlikely | Unlikely |
| Arc State (SE) | Unlikely | Unlikely | Likely (Census Jan 2025) | Unlikely | Unlikely |
| Arc State (ST) | Unlikely (not in training list) | **Confirmed** (Replogle-Nadig) | N/A (ST not appl. Dim D) | N/A (ST not appl. Dim E) | N/A (ST not appl. Dim E) |

**Supplementary Table 6.**
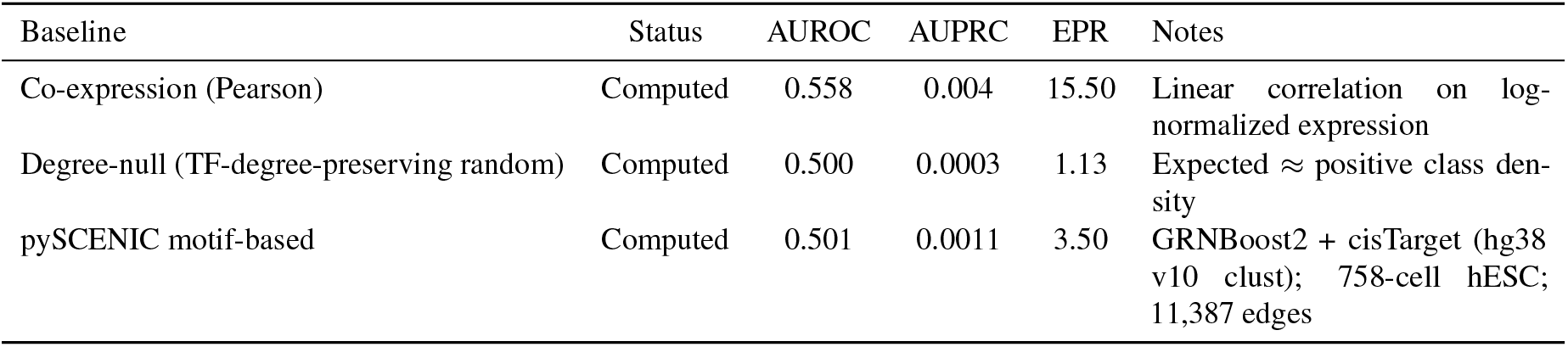
GRN baseline status and passing rule components. The pre-registered passing rule for Dimension C (GRN inference) required AUPRC on directed TF-to-target edges to exceed all three baselines: co-expression, degree-null, and pySCENIC. All three are computed. pySCENIC (AUPRC 0.0011) sits between the degree-null floor (0.0003) and co-expression (0.004). scGPT is the only foundation model to exceed pySCENIC (AUPRC 0.003 vs 0.0011) but still falls short of co-expression (0.003 vs 0.004). No foundation model clears all three baselines.

**Supplementary Table 7.** Virtual Cell Level assignments per model and per dimension. Each cell records whether the foundation model exceeds the trivial baseline (mean-prediction or no-change baseline; first value) and whether it exceeds the strongest non-foundation-model baseline (second value) on the given dimension. ✓indicates the condition is met, x indicates it is not; DNR, not executed; N/A, structural architectural incompatibility. The final column reports the assigned VC Level following the pre-registered mapping from Noutahi et al. [9] as operationalized by VCBench: Level 1 requires exceeding the trivial baseline on ≥1 dimension; Level 2 requires exceeding the strongest non-foundation-model baseline on ≥1 dimension; Level 3 requires exceeding the strongest non-foundation-model baseline on ≥3 dimensions. The Dimension C “beats strongest baseline” criterion requires exceeding all three computed baselines (co-expression, degree-null, and pySCENIC); see Supplementary Table 6. Key observations: (i) all five foundation models achieve Level 1 (Geneformer, scGPT, UCE, Transcript-Former on multiple dimensions; Arc State on Dimension A via no-change baseline only); (ii) one model achieves Level 2 (TranscriptFormer on Dimension D); (iii) no model achieves Level 3. TranscriptFormer is the closest to Level 3, clearing the strongest baseline on Dimension D while scoring below baseline on A, B (under matched class sets), and E and reported N/A on C (see Methodological notes). These assignments are visualized in Figure 8.

| Model | Dim A (triv./str.) | Dim B | Dim C | Dim D | Dim E | Trivial cnt | Strong. cnt | VC Lvl |
| --- | --- | --- | --- | --- | --- | --- | --- | --- |
| Geneformer V2-316M | ✓ / x | ✓ / x | ✓ / x | ✓ / x | x / x | 4 | 0 | 1 |
| scGPT (fine-tuned) | ✓ / x | ✓ / x | ✓ / x | ✓ / x | x / x | 4 | 0 | 1 |
| UCE 33-layer | N/A | ✓ / x | N/A | ✓ / x | ✓ / x | 3 | 0 | 1 |
| TranscriptFormer | x / x | ✓ / x | N/A | ✓ / ✓ | ✓ / x | 3 | 1 | 2 |
| Arc State | ✓ / x | N/A | N/A | DNR | DNR | 1 | 0 | 1 |

**Supplementary Table 8.** Spread-error correlation probe: LFC spread versus prediction error. Spearman rank correlation (*ρ*) between per-perturbation LFC spread (variance of predicted log-fold changes across 5,045 genes) and per-perturbation MAE on Norman CRISPRa (GEARS standard split, 107 test perturbations). Positive *ρ* indicates the model assigns higher spread to harder predictions (spread-error correlation signal present); *ρ* ≈ 0 indicates no signal; negative *ρ* indicates inverse-spread behavior. Neither tested model achieves statistical significance (*p >* 0.17), indicating that LFC spread carries no information about prediction quality. This is a necessary but not sufficient negative result for epistemic calibration: true calibration would require a distributional output, an ensemble, MC dropout, or a conformal wrapper, none of which the tested models provide. UCE is N/A (no expression-level output). TranscriptFormer was not evaluated because its autoregressive architecture requires a full forward pass per cell and per perturbation condition, with projected inference time exceeding 5 hours for 245 perturbation conditions on the Norman dataset. Arc State lacks a native perturbation prediction API in its public release and would require training from scratch, which was out of scope for this supplementary analysis.

| Model | Spearman $\rho$ | $p$-value | $n$ | Entropy<br>(mean ± std) | MAE<br>(mean ± std) | Interpretation |
| --- | --- | --- | --- | --- | --- | --- |
| Geneformer V2-316M | -0.119 | 0.225 | 106 | 0.0012 ± 0.0005 | 0.0185 ± 0.0088 | No detectable correlation (n.s., $p = 0.225$) |
| scGPT (fine-tuned) | +0.131 | 0.177 | 107 | 0.0019 ± 0.0034 | 0.0203 ± 0.0073 | No detectable correlation (n.s., $p = 0.177$) |
| UCE 33-layer | — | — | — | — | — | No expression-level output |
| TranscriptFormer | — | — | — | — | — | Autoregressive inference prohibitively slow |
| Arc State | DNR | DNR | — | — | — | No perturbation prediction API |

### Supplementary Note 1. VCBench Contamination Reporting Schema v1

A minimal machine-checkable standard for single-cell foundation-model training-data provenance, released alongside the VCBench benchmark. The goal is to make it possible for a benchmark author to run `vcbench-check-contamination model_weights_dir/ eval_dataset.h5ad` and receive a definitive `confirmed/likely/unlikely/unknown` verdict in under one second, replacing the platform-inference audit VCBench currently applies (Supplementary Tables 4 and 5) with direct overlap detection.

#### Scope and motivation

Single-cell foundation-model papers (Geneformer, scGPT, UCE, TranscriptFormer, Arc State) currently publish training-corpus summaries at the dataset level (for example, “CELLxGENE Census, Jan 2025 build”) but not at the cell level. This makes exhaustive contamination detection between pretraining and evaluation datasets impossible. As a result, a benchmark like VCBench can only assert `likely` or `unlikely` based on platform inclusion/exclusion rules. NLP addressed the analogous problem with n-gram overlap detection and canonical decontamination protocols. Single-cell biology does not yet have an equivalent standard.

The schema described here is minimal by design: one manifest file per released model, one metadata flag per evaluated cell. It does not require models to withhold data they did not withhold, and it does not require retraining. It requires disclosure only.

#### Component 1. Cell-barcode MD5 manifest

Every model released alongside a pretraining corpus publishes a file named `training_cells.md5.txt` or `training_cells.md5.parquet` at the root of the release. The file contains one row per cell used in pretraining, with columns barcode_md5 (32-hex-char MD5 hash of the cell barcode), `source_accession` (GEO, SRA BioProject, Figshare DOI, or CELLxGENE Census dataset ID), and `corpus_version` (e.g. `census-2025-01-15)`. The MD5 hash is computed on the raw barcode string (no whitespace, no source-ID prefix), encoded UTF-8. The hash makes privacy-safe publication straightforward while still permitting exact intersection tests: a benchmark can MD5-hash its own evaluation-set barcodes and intersect against the published manifest.

#### Component 2. AnnData .obs split flag

Models that are released with held-out evaluation data (any paper that reports a held-out metric) include an.`obs` column named `vcbench_pretrain_split` with one of the following values per cell: `train` (used in pretraining), `val` (held out for validation during pretraining), `test` (held out entirely from pretraining, safe for downstream benchmarking), or `unknown` (provenance not recorded). The `unknown` value is permitted but must not exceed 10% of cells in any released dataset; otherwise the release does not satisfy the schema.

#### Component 3. Accession-level manifest

A human-readable file named `pretraining_manifest.yaml` in the release root listing every dataset used for pretraining at the accession level. The exclusions list uses canonical reasons from a short controlled vocabulary `(cell_culture, perturbation_assay, non_human, insufficient_metadata, cohort_restricted, other)`. Accessions that were considered and rejected must be listed; silent exclusions do not satisfy the schema. Example:

~~~
corpus_version: “census-2025-01-15”
model_name: “example-foundation-model-v2”
sources:
 - accession: “GSE194122”
  platform: “GEO”
  included: true
  cells_used: 66175
  inclusion_criteria: “bone marrow CITE-seq, all four sites”
  exclusions: []
  notes: “RNA component only; ADT component not used”
 - accession: “10.25452/figshare.plus.20029387”
 platform: “Figshare+”
 included: false
 cells_used: 0
 inclusion_criteria: null
 exclusions: [“cell_culture”, “perturbation_assay”]
 notes: “Excluded per Census cell-culture filter”
~~~

#### Validator behavior

A compliant validator `(vcbench-check-contamination)` takes a model release directory and one or more evaluation datasets and returns, for each (model, evaluation-dataset) pair: confirmed (non-empty MD5 intersection); `likely` (no MD5 intersection, but evaluation-dataset accession is listed as `included); unlikely` (no MD5 intersection, and evaluation-dataset accession is listed as `excluded)`; or `unknown` (no MD5 intersection and evaluation-dataset accession not mentioned in manifest).

#### Reference implementation

A reference Python implementation lives at `tools/vcbench_contamination_check/` in the VCBench repository. The validator core is approximately 200 lines of Python; dependencies are `pyyaml, pandas`, and `anndata`. The stub below illustrates the core logic; production implementation adds CLI argument parsing, logging, and progress reporting.

~~~
from pathlib import Path
import hashlib, yaml
import pandas as pd
import anndata as ad
def check_contamination(model_release_dir: Path, eval_dataset: Path) -> dict:
  “““Return a contamination verdict for one model-evaluation pair.”““
  model_release_dir = Path(model_release_dir)
  eval_dataset = Path(eval_dataset)
  # 1. Confirm schema compliance
  md5_file = model_release_dir / “training_cells.md5.parquet”
  manifest_file = model_release_dir / “pretraining_manifest.yaml”
  missing = [f for f in (md5_file, manifest_file) if not f.exists()]
  if missing:
    return {“verdict”: “schema_incomplete”,
              “missing”: [str(f) for f in missing]}
  # 2. Load pretraining MD5 set
  md5_df = pd.read_parquet(md5_file)
  pretrain_md5 = set(md5_df[“barcode_md5”])
  # 3. Hash evaluation-dataset barcodes
  adata = ad.read_h5ad(eval_dataset)
  eval_barcodes = adata.obs_names.astype(str)
  eval_md5 = {hashlib.md5(b.encode(“utf-8”)).hexdigest()
         for b in eval_barcodes}
   intersection = pretrain_md5 & eval_md5
   if intersection:
      return {“verdict”: “confirmed”,
            “overlap_cells”: len(intersection),
            “overlap_fraction”: len(intersection) / len(eval_md5)}
  # 4. No MD5 overlap; consult accession manifest
  manifest = yaml.safe_load(manifest_file.read_text())
  eval_accession = adata.uns.get(“source_accession”)
  if eval_accession is None:
     return {“verdict”: “unknown”,
           “reason”: “eval dataset has no source_accession in .uns”}
  matched = [s for s in manifest[“sources”]
        if s[“accession”] == eval_accession]
  if not matched:
    return {“verdict”: “unknown”,
       “reason”: f”{eval_accession} not listed in manifest”}
  entry = matched[0]
  if entry[“included”]:
    return {“verdict”: “likely”,
       “rationale”: entry.get(“notes”, ““)}
  return {“verdict”: “unlikely”,
       “exclusions”: entry.get(“exclusions”, []),
       “rationale”: entry.get(“notes”, ““)}
~~~

#### Adoption pathway

The schema is deliberately sized so that adoption costs are dominated by authorship, not computation. MD5-hashing a pretraining corpus of 100 million cell barcodes takes under ten minutes on a single CPU. Writing the accession-level manifest takes an hour of an author’s time for a typical foundation-model paper. The `vcbench_pretrain_split` column is one line of `adata.obs` assignment.

Future schema versions may add cryptographic commitment hashes that permit third-party verification without exposing barcode-level data (v2); canonical per-dataset hashes for public datasets, so that “same dataset, different versions” can be distinguished (v3); and integration with CELLxGENE Census versioning to permit automated inclusion/exclusion lookups (v3).

#### Availability

Specification and reference implementation released under CC-BY-4.0 (spec) and MIT (code) within the VCBench repository at github.com/AppliedScientific/VCBench (release tag `v1.0.0,` commit `079355d`). The schema is versioned separately from the benchmark code so that adopting models can cite a stable specification.

### Supplementary Note 2. Pre-submission methodological audit and operational caveats

This note collects the audit narratives and operational caveats referenced from main Methods and Discussion. Items are grouped by audit subject; each subsection records the issue, scope, resolution, and any remaining limitations.

#### S2.1. TranscriptFormer Dimension C scoring path

A pre-submission code-level audit of `src/models/run_transcriptformer_grn.py:: step1_embedding_coexpression()` (lines 34–92) revealed that the function did not instantiate the TranscriptFormer model. The script’s docstring (lines 4–12) described the intended procedure: extract baseline cell embeddings, then for each transcription factor zero out that gene’s counts, re-embed, and score TF target edges by the correlation between target-gene expression and the embedding shift. The implementation instead computed `abs(Pearson correlation)` on raw expression in the BEELINE hESC corpus (lines 59–66, 82). No model checkpoint was loaded; no forward pass occurred. The originally reported TranscriptFormer Dimension C numbers (AUROC 0.510, AUPRC 0.001, EPR 4.81) were therefore co-expression scores under a different normalization rather than TranscriptFormer-derived scores.

The audit was conducted as part of the same pre-submission methodological pass that surfaced the GRN passing-rule amendment and the train-test leak in our Arc State evaluation documented in main Methods. Independent verification of the Geneformer V2-316M scoring path `(run_geneformer_grn.py:` layer-13 attention aggregation across the hESC corpus) and the scGPT scoring path `(run_scgpt_grn.py:` cosine similarity on learned gene embeddings from the released checkpoint) confirmed both matched their Methods descriptions.

In place of attempting to re-implement and re-validate the masked-probability procedure under pre-submission time pressure, with attendant risk of introducing new defects, TranscriptFormer is reported N/A on Dimension C. This treatment is consistent with the existing N/A annotations for UCE and Arc State on Dimension C (architectural rather than implementation gaps). The TranscriptFormer Dimension D headline result uses a different code path that was verified to operate correctly. TranscriptFormer’s VC Level assignment (Level 2 via Dimension D) is unchanged because the original Dimension C numbers did not satisfy the strongest-baseline criterion under either the original or the amended passing rule.

#### S2.2. Dimension A pipeline reconciliation

A pre-submission audit of `results/tables/table2_with_baselines.csv` against the `cell_eval` outputs in `results/dim_a/*/cell_eval_results.csv` identified a staleness drift on four cells of Table 2: Geneformer V2-316M Dimension A PRR (0.5772 ⟶ 0.6267) and DES (0.8703 ⟶ 0.8778), and scGPT Dimension A PRR (0.4944 ⟶ 0.5025) and DES (0.8640 ⟶ 0.8439). The drift was traced to a pipeline refresh dated 2026-04-17 whose underlying outputs were never reflected in the version of Table 2 carried into the original draft. TranscriptFormer Dimension A numbers were unaffected because TranscriptFormer was re-evaluated under the refreshed pipeline in a 2026-04-26 bundle. Dimensions B, C, D, and E were verified consistent with current pipeline outputs across all evaluated models. The four affected cells, plus their associated composite scores, were updated to canonical pipeline values.

Subsequent inspection of the partition decomposition file `subset_analysis.json` revealed that it was produced by the same pre-refresh pipeline (file mtime 2026-04-15, matching the stale Table 2 values exactly). The partition decomposition was therefore recomputed against the canonical pipeline using a fresh scGPT fine-tune on Norman (15 epochs, AMP, seed 42; vanilla MHA fallback used because flash-attention does not build against the env’s torch+CUDA combination). The recomputed novel-subset PRR for scGPT moved from 0.4194 to 0.4196, an effective non-shift; the larger pipeline-refresh delta was concentrated in the shared subset (Δ = +0.012). All v2 numerical updates rest on canonical pipeline outputs; the qualitative claims are unaffected. Fresh scGPT predictions and the partition file are released alongside the benchmark code to support reproducibility.

#### S2.3. Dimension B native-vs-common-set protocol comparison

Initial Dimension B numbers were computed under a native-label protocol in which each foundation model was scored on the cell-type vocabulary surviving its own tokenizer and filter. Under this protocol Geneformer’s macro F1 (0.181 aggregate) appeared to exceed the PCA+kNN baseline (0.166 aggregate), and TranscriptFormer’s weighted F1 on lung+liver (0.347) appeared to exceed the matched baseline value, supporting two Level-2 attributions. A pre-submission audit identified a class-count confound: macro F1 averaged across cell-type classes is mechanically lower when computed across more classes, and the foundation-model evaluations admitted fewer classes per tissue than the baseline did because of tokenizer-driven cell filtering. The Geneformer ranking in particular reflected scoring against a smaller cell-type vocabulary rather than superior cross-species transfer.

To remove the confound, both foundation models and baselines were re-evaluated under a common-label-set protocol that restricts every method per tissue to the intersection of cell-type vocabularies admitted by all evaluated models and the baseline. Under this protocol PCA+kNN aggregate macro F1 is 0.497, exceeding Geneformer common-set 0.171, scGPT 0.123, UCE 0.379, and TranscriptFormer common-set 0.351 (lung+liver subset). Per-tissue, PCA+kNN exceeds every foundation model on lung, heart, kidney, and brain; on liver, TranscriptFormer (macro F1 0.495) marginally exceeds PCA+kNN (0.446) by a relative margin of 11%. The lung+liver aggregate (0.456 baseline vs 0.351 TF) averages over this per-tissue inversion. Both Level-2 attributions previously assigned on Dimension B are removed under the common-label-set protocol; TranscriptFormer’s Dimension D attribution is unaffected because Dimension D does not use a class-based metric. The TranscriptFormer common-set evaluation was completed on lung and liver only, the two tissues where TranscriptFormer holds species embeddings; full-tissue common-set evaluation is scoped to a future revision and would require re-extraction with cell-type labels persisted to embedding metadata. Both species’ embeddings were recovered for the existing two-tissue analysis by leveraging row-order preservation in source AnnData files, requiring no GPU re-run.

#### S2.4. Operational caveat on the contamination audit

The contamination audit identifies known overlaps between training corpora and evaluation datasets but cannot be exhaustive. No evaluated model publishes a complete cell-level training manifest at the GEO accession level; classification therefore relies on platform-level inclusion and exclusion criteria (CELLxGENE Census exclusions, Genecorpus-30M cell-line exclusions, Arc State documented training datasets). NLP-derived contamination detection methods (n-gram overlap, membership inference) have no direct single-cell analog. The proposed Contamination Reporting Schema v1 (Supplementary Note 1) is the proposed remedy.

#### S2.5. Operational caveat on the spread-error correlation probe

The spread-error correlation probe (Supplementary Table 8) covers two of five foundation models on Norman: Geneformer V2-316M and scGPT. TranscriptFormer was not evaluated due to prohibitive autoregressive inference time (projected *>* 5 hours per perturbation condition); Arc State lacks a public perturbation prediction API in its released configuration. The consistent non-significance across both tested models (Spearman |*ρ* | *<* 0.15, *p >* 0.17) supports the empirical claim that LFC spread does not correlate with prediction error in current foundation models, but coverage across additional models and datasets, together with a direct calibration experiment (heteroscedastic output head, ensemble, or conformal wrapper) on at least one model, would strengthen the conclusion. The probe is necessary but not sufficient evidence for absence of epistemic calibration.

### Supplementary Note 3. Detailed per-dimension evaluation protocols

This note collects the per-model fine-tuning hyperparameters, library versions, and protocol specifics underlying each scored dimension.

References to Table 2 and figure numbers follow the preprint numbering.

#### Dimension A (perturbation prediction)

Predictions are evaluated using a metric suite derived from Arc Cell-Eval [7, 8]: Perturbation Response Recovery (PRR), Direction Score (DES; sign-agreement on the top-20 most-perturbed genes per perturbation, see Eq. 3), and Mean Absolute Error (MAE) on pseudobulk profiles. PRR is computed by VCBench’s evaluation pipeline as the mean per-perturbation Pearson correlation between predicted and observed log-fold-change vectors relative to control. Cell-eval (PyPI release dated 2026-04-09) is invoked through `cell_eval.MetricsEvaluator` as the entry point; the FM evaluation path falls through to a custom implementation that computes both PRR (per-perturbation Pearson on Δ-expression) and DES (sign-agreement on the top-20 Δ-expression genes) directly on pseudo-bulk averages rather than the canonical Cell-Eval rank-based discrimination and gene-level Wilcoxon procedures. We adopt these renamed metrics in lieu of Cell-Eval’s canonical PDS (a ranking-based discrimination metric on a closed candidate-perturbation pool) and canonical DES (a per-cell Wilcoxon-based score) because the held-out single-perturbation evaluation setting and pseudo-bulk pipeline used here do not match those canonical procedures. The additive-baseline DES value reported in canonical Cell-Eval mode (0.5665) is therefore not directly comparable to the FM DES values reported in this manuscript, paralleling the PRR/PDS distinction. Inputs were detected as log1p-normalized and passed through without additional transformation.

#### Per-model perturbation prediction protocols

Geneformer V2-316M was fine-tuned as a `BertForSequenceClassification` with one class per Norman perturbation (247 classes total) for 3 epochs at learning rate 5 ×10^−5^, per-device batch size 12, with the bottom 2 transformer layers frozen, 100 warmup steps, weight decay 0.01, and the AdamW optimizer. Post-fine-tuning, cell embeddings were extracted from the last hidden layer (`emb_layer=0,` following Ahlmann-Eltze et al., 2025). Perturbation was simulated by token deletion: perturbed gene tokens were removed from the input rank-value sequence and the model was re-embedded. A ridge regression decoder (α = 1.0) was trained on control-plus-training-perturbation embeddings mapping to expression vectors, then applied to test-perturbation embeddings to produce predicted post-perturbation expression. scGPT was fine-tuned via the `TransformerGenerator` architecture with masked MSE loss for up to 15 epochs at learning rate 1 ×10^−4^, batch size 16, StepLR schedule (*γ* = 0.9), maximum sequence length 1,536, and early stopping on validation loss with patience 10. Inference used the native `pred_perturb` API with the fine-tuned checkpoint. TranscriptFormer was evaluated zero-shot with a ridge regression decoder (α = 1.0). Inference configuration was built in-process via OmegaConf with `pretrained_embedding=True` and `output_keys=[“embeddings”]`; no on-disk YAML was used. Perturbation was simulated by gene zeroing: for each test perturbation with target genes *G*, input expression values were set to zero at the corresponding gene indices before re-embedding. The ridge decoder was fit on train-set embeddings (control and training perturbations, with test perturbation IDs held out per the `norman_ gears_split.toml` configuration) and applied to test perturbations. Predictions were clipped to the non-negative range [0, GT_max] to prevent ridge-regression extrapolation to implausible expression values.

Arc State was trained from scratch on Norman using the configuration in `configs/norman_gears_split.toml`, which enumerates 138 training perturbations plus the non-targeting control and 107 held-out test perturbations with zero overlap (the canonical GEARS simulation split, seed 1). The training hyperparameters match the published Arc State Norman recipe [7] `(cell_set_len = 512`, batch size 8, transformer backbone with 8 hidden layers, AdamW optimizer at learning rate 1 × 10^−4^, 40,000 steps); the only deviation from the public release is the corrected GEARS split documented in Methodological note: train-test leak in our Arc State evaluation. Training loss converged from 2.94 to 0.027. Validation loss reached a minimum of 0.263 at ~step 30,000 and rose to ≈0.40 at step 39,999 (overfitting signature on the 138-perturbation training set). The PRR we initially obtained for Arc State on Norman was inflated by a train-test leak in our earlier configuration `(norman_fewshot.toml)` in which a cell-type filter string we wrote matched zero rows, causing all 107 test perturbations to remain in the training pool. The leaked configuration produces PRR 0.949 on the GEARS test set when scored through the corrected evaluator (because the model is being scored on data it saw during training); the corrected configuration produces PRR 0.402 on the same test set. Three independent signatures confirm the overlap: under the leaked configuration the validation DataLoader is empty (no `val_*` columns are ever logged during training), `state tx predict` structurally crashes on the leaked checkpoint without an explicit split override, and the leaked checkpoint’s high PRR is achieved on perturbations the model trained on. We notified Arc Institute of the silent-filter behavior prior to preprint posting; the corrected retrained result (PRR 0.402) is the one reported in all tables, and the corrected checkpoint is released alongside the benchmark code.

#### Evaluator anchor convention

The vcbench evaluator computes per-perturbation Pearson correlation against a shared real-control anchor: 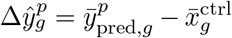 and 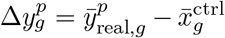, treating the control mean as a property of the data rather than the model. Cell-Eval’s `pearson_delta` defaults to a per-model anchor convention in which the predicted control mean is subtracted from predicted perturbation means. The two conventions agree exactly when the model’s predicted control matches the real control per-gene; they diverge when it does not. For Arc State on Norman the per-gene maximum absolute control divergence was 0.066, producing a PRR gap of 0.005 under the corrected configuration (vcbench 0.402 vs Cell-Eval 0.408) and 0.015 under the leaked configuration (vcbench 0.949 vs Cell-Eval 0.964). VCBench adopts the shared real-control anchor as canonical for Virtual Cell Level decisions because it does not allow models to absorb baseline drift through their predicted control, which is particularly important when evaluating leaked configurations where memorized baselines would otherwise inflate apparent recovery. Both conventions are exposed via the `control_anchor` parameter of `evaluate_dim_a` (Eq. 1); under matched conventions the vcbench evaluator reproduces Cell-Eval to numerical precision (max absolute per-perturbation difference *<* 10^−6^ on real Arc State predictions, *<* 10^−9^ on synthetic data), with the equivalence locked by a unit test.

#### Matched-regime experiment (Geneformer ZS+D vs FT+D)

To distinguish regime from representation in the Dim A cross-model comparison, the same Geneformer V2-316M checkpoint was evaluated in a zero-shot-with-decoder (ZS+D) regime on the Norman task, alongside the fine-tuned-with-decoder (FT+D) regime described above. The ZS+D protocol uses the pretrained Geneformer V2-316M model without fine-tuning. Cell embeddings were extracted from the last hidden layer at `emb_layer=0`. Perturbation was simulated by token deletion (identical to the FT+D protocol): perturbed gene tokens were removed from the input rank-value sequence and the model was re-embedded. A ridge regression decoder (− = 1.0) was fit on per-perturbation embedding means computed across control-plus-training-perturbation cells, then applied to test-perturbation embedding means to produce predicted post-perturbation expression. The ZS+D and FT+D regimes share all components (decoder family, perturbation simulation, evaluation set) except the fine-tuning step, isolating regime as the experimental variable.

For the Dimension A evaluation, baselines are: the Ahlmann-Eltze additive model (*y*(*A* + *B*) = ctrl + LFC(*A*) + LFC(*B*)), a mean-prediction baseline (mean of training perturbation expression), and a no-change baseline (control expression). The additive baseline is evaluable only on perturbations whose single-gene components appear in the training set (71 of 107 on Norman); head-to-head comparison with foundation models is reported on this shared 71-perturbation subset in Results.

#### Dimension B (cross-species)

Cell embeddings are extracted for human and mouse cells per tissue. A k-nearest-neighbors classifier (*k* = 5) trained on human embeddings predicts cell type labels for mouse cells. Metrics: macro-averaged F1 and weighted F1. Baseline: PCA (50 components) on log-normalized ortholog-mapped expression followed by the same kNN classifier. Ortholog mapping uses `pybiomart` against live Ensembl BioMart (release approximately 113 at time of execution, April 2026) with a three-host fallback chain (www.ensembl.org, useast.ensembl.org, asia.ensembl.org), filtering on `homology_type == “ortholog_one2one”` and `confidence = 1.` This yields 15,847 high-confidence one-to-one human–mouse orthologs. The ortholog map is cached deterministically at `data/processed/ortholog_maps.pkl`; a static JSON backup (`data/static/ensembl_symbol_map.json`) is committed to the repository for reproducibility.

Tissue coverage varies by model: Geneformer and scGPT were evaluated on all five tissues (lung, liver, heart, kidney, brain); TranscriptFormer was evaluated on lung and liver only; UCE was evaluated on heart and brain only, due to species-vocabulary compatibility and compute budget. Matched-tissue head-to-head comparisons are reported separately to avoid aggregate-denominator confounds.

#### Dimension C (GRN inference)

Regulatory edges are extracted from model internals and evaluated against BEELINE [17] and TRRUST v2 [18] ground truth using AUROC, AUPRC, and Early Precision Ratio (EPR) on directed transcription factor-to-target edges. For Geneformer, attention weights were aggregated at layer 13 across all cells in the hESC corpus; edge scores are the mean attention from TF token positions to target token positions. For scGPT, edge scores are cosine similarity on learned gene embeddings. The scGPT Dimension C protocol uses static token-embedding cosine similarity from the released checkpoint, without task-specific fine-tuning or cell-conditioning; it is the most parsimonious of the three Dimension C protocols, which contributes to scGPT’s modest performance on this dimension. For TranscriptFormer, the Methods originally described conditional token probability shifts via masking individual TF tokens and measuring log-probability changes at other gene positions; pre-submission code audit revealed that the available codebase did not implement this procedure and instead fell back to absolute Pearson correlation on raw expression in the hESC corpus, yielding co-expression-equivalent rather than model-derived scores. The originally reported TranscriptFormer Dimension C numbers were retracted on this basis. TranscriptFormer is reported N/A on Dimension C in all tables and figures; see “Methodological note: TranscriptFormer Dimension C.” UCE is structurally N/A (input gene tokens are static ESM-2 protein embeddings that encode sequence similarity rather than learned regulatory relationships).

Baselines: Pearson co-expression correlation on log-normalized expression, a degree-null model that samples random TF-target edges preserving the empirical TF out-degree distribution, and a pySCENIC motif-based baseline [38]. pySCENIC was run on the 758-cell BEELINE hESC corpus using GRN-Boost2 for co-expression module inference followed by cisTarget motif pruning (hg38 v10 clustered ranking databases, 10 kb upstream/downstream of TSS), producing 11,387 regulatory edges in 78 minutes on an 8-vCPU / 15 GB CPU-only server (no GPU required). All models and baselines are scored on the same TF universe: the intersection of TRRUST v2 transcription factors with each model’s expressed gene vocabulary, computed once per model and applied uniformly across that model’s predictions and the matched baseline computation, ensuring apples-to-apples comparison via the canonical `evaluate_and_save_grn()` function. Where a model’s tokenizer admits a smaller subset of TRRUST TFs than another model, the per-model TF universe differs accordingly; cross-model comparison is therefore a comparison across slightly different TF subsets (typically 795 TFs versus 800), an asymmetry that is preserved by the uniform within-model evaluation but disclosed here for completeness. The TRRUST v2 database contains 8,444 published TF-target pairs across 800 TFs; the evaluated subset is smaller after restricting to the shared gene vocabulary (typically 8,400 edges across 795 TFs, varying slightly by model). The pre-registered passing rule requires AUPRC to exceed all three baselines (co-expression, degree-null, and pySCENIC); all three are computed and reported.

#### Dimension D (cross-modal prediction)

Cell embeddings are extracted from RNA features only on the training split (three sites). A ridge regression (*α* = 1.0) maps embeddings to 134 protein expression values. Predictions are evaluated on the held-out fourth site using per-protein Pearson correlation (mean across 134 proteins) and RMSE. All five foundation models participate through the embedding probe protocol. Baselines: mean celltype (predict the mean protein expression of each training-site cell’s predicted cell type), PCA (50 components) plus ridge regression, and scLinear [39]. The mean celltype baseline establishes the performance floor below which a model is providing less information than cell type identity alone.

#### Dimension E (temporal ordering)

Cell embeddings are extracted for all cells in each time-course dataset. Diffusion pseudotime [40] was chosen over alternatives (RNA velocity, Waddington-OT) because it operates on a generic embedding space without requiring spliced/unspliced count decomposition or explicit cost functions, making it applicable to all five foundation models on equal footing. For methodological consistency with the PCA+DPT baseline, all foundation model embeddings are first reduced to 50 principal components before downstream analysis (native dimensions: Geneformer 1,152, scGPT 512, UCE 1,280, TranscriptFormer 2,048; PCA variance retained: 0.72-0.99 across models and datasets). A kNN graph (*k* = 15) is built on the PCA-reduced embeddings, followed by diffusion pseudotime [40] computation with root cell from the earliest timepoint. Temporal ordering is evaluated by Kendall’s tau-b between DPT and true collection time. Discrete timepoint prediction is evaluated by kNN balanced accuracy (*k* = 15, 5-fold cross-validation). Baseline: PCA (50 components) followed by DPT on log-normalized HVG-selected counts.

For LARRY (mouse data), human-vocabulary models (Geneformer, scGPT) use ortholog mapping to human Ensembl IDs before embedding extraction, matching the cross-species protocol; Arc State is N/A on LARRY because its tokenizer has no mouse-gene mapping logic. TranscriptFormer’s embeddings on the full 49,008-cell Weinreb dataset produced a degenerate graph Laplacian during diffusion map eigendecomposition: ARPACK’s sparse Lanczos iteration failed to converge `(ArpackError -9999)` because TranscriptFormer embeds near-duplicate cells with zero pairwise distance, producing disconnected kNN components and many indistinguishable near-zero eigenvalues. The pathology is size-sensitive: 5,000-cell subsamples converge in under 10 seconds, while the full 49,008-cell graph does not converge within ARPACK’s maximum iteration budget. TranscriptFormer’s Weinreb result is therefore reported as a bootstrap estimate (mean and standard deviation over 10 random 5,000-cell subsamples; *τ* = 0.041 *±* 0.078), flagged in Table 2 with a dagger.

Per-model Dimension E aggregates in Table 2 are the unweighted arithmetic mean of the per-dataset Kendall’s *τ* -b values (sci-fate and Weinreb), giving each dataset equal weight regardless of cell count. This convention preserves the dataset-level signal (in particular, the scGPT temporal-inversion finding on Weinreb) which would be diluted under a cell-count-weighted aggregation given Weinreb’s larger sample. Per-dataset breakdowns are reported in Supplementary Table 3.

#### Spread-error correlation probe (epistemic calibration)

For each model’s perturbation predictions on Norman (GEARS standard split, 107 test perturbations, 5,045 genes), LFC spread is computed as the variance of predicted log-fold changes across genes for each perturbation. Spearman rank correlation between per-perturbation LFC spread and per-perturbation MAE is reported. Positive correlation indicates the model assigns higher spread to harder predictions (spread-error correlation signal present); zero correlation indicates no signal; negative correlation indicates inverse-spread behavior. Geneformer (fine-tuned, 5.7 h on H200 GPU) and scGPT (fine-tuned, 2.75 h on H200 GPU) were evaluated. TranscriptFormer was not evaluated because its autoregressive architecture requires a full forward pass per cell and per perturbation condition, with projected inference time exceeding 5 hours for the 245 Norman perturbation conditions alone. Arc State lacks a native perturbation prediction API in its public release. UCE is structurally N/A (no expression-level output). Results are reported in Supplementary Table 8.

This probe measures whether the spread of a point prediction across output dimensions correlates with per-perturbation error. It is a weaker and more indirect signal than true calibration of a predictive distribution (which would require a distributional output, an ensemble, MC dropout, or a conformal wrapper). A failure to detect spread-error correlation is a necessary but not sufficient negative result for epistemic calibration, reported here as the minimal prerequisite the active-learning loops envisioned by Bunne et al. [2] and Noutahi et al. [9] would demand.

#### Contamination audit

For each model-dataset pair, VCBench classifies contamination risk as *confirmed* (dataset is documented in training corpus), *likely* (dataset meets all inclusion criteria of training data source and no exclusion is documented), *unlikely* (dataset fails documented exclusion criteria of training source), or *unknown* (training manifest is insufficient to determine). Classification is based on published training data documentation: paper supplementary materials, data repository manifests, and platform schema (CELLxGENE Census inclusion/exclusion criteria). CELLxGENE Census excludes observations with `tissue_type “cell culture”` and perturbation-based assays. Geneformer’s Genecorpus-30M excludes immortalized cell lines and malignant cells. Arc State’s transition model training datasets are enumerated in Adduri et al. [7]. Where training manifests are incomplete (scGPT, UCE, TranscriptFormer), risk is assessed from platform-level inclusion criteria. Results are reported in Supplementary Table 5.

#### VC Level mapping

VC Level definitions, threshold-based passing criteria, and per-dimension trivial and strongest baselines are documented in Methods §VC Level mapping (body) and visualized in Figure 1d.

## Notes

https://github.com/AppliedScientific/VCBench

https://huggingface.co/collections/appliedscientific/vcbench-v100-single-cell-foundation-model-benchmark

### Competing Interest Statement

The authors have declared no competing interest.

